# The Dream Catcher experiment: Blinded analyses disconfirm markers of dreaming consciousness in EEG spectral power

**DOI:** 10.1101/643593

**Authors:** William Wong, Valdas Noreika, Levente Móró, Antti Revonsuo, Jennifer Windt, Katja Valli, Naotsugu Tsuchiya

**Affiliations:** School of Psychological Sciences and Turner Institute for Brain and Mental Health, Monash University, Melbourne, Victoria, Australia; Department of Psychology, University of Cambridge, Cambridge, United Kingdom; Department of Psychology, and Turku Brain and Mind Center, University of Turku, Turku, Finland; Department of Cognitive Neuroscience and Philosophy, School of Bioscience, University of Skövde, Skövde, Sweden; Philosophy Department, Monash University, Clayton, Victoria, Australia; Department of Dynamic Brain Imaging, Advanced Telecommunications Research Institute International, Seika, Kyoto Prefecture, Japan; Center for Information and Neural Networks, National Institute of Information and Communications Technology, Suita, Osaka Prefecture, Japan

**Keywords:** NREM sleep, dreams, unconsciousness, EEG correlates, unsupervised machine learning

## Abstract

The Dream Catcher test defines the criteria for a genuine discovery of the neural constituents of phenomenal consciousness. Passing the test implies that some patterns of purely brain-based data directly correspond to the subjective features of phenomenal experience, which would help to bridge the explanatory gap between consciousness and brain. Here, we conducted the Dream Catcher test for the first time in a graded and simplified form, capturing its core idea. The experiment involved a Data Team, who measured participants’ brain activity during sleep and collected dream reports, and a blinded Analysis Team, who was challenged to predict better than chance, based solely on brain measurements, whether or not a participant had a dream experience. Using a serial-awakening paradigm, the Data Team prepared 54 one-minute polysomnograms of NREM sleep—27 of dreamful sleep (3 from each of the 9 participants) and 27 of dreamless sleep—redacting from them all associated participant and dream information. The Analysis Team attempted to classify each recording as either dreamless or dreamful using an unsupervised machine learning classifier, based on hypothesis-driven, extracted features of EEG spectral power and electrode location. The procedure was repeated over five iterations with a gradual removal of blindness. At no level of blindness did the Analysis Team perform significantly better than chance, suggesting that EEG spectral power does not carry any signatures of phenomenal consciousness. Furthermore, we demonstrate an outright failure to replicate key findings of recently reported correlates of dreaming consciousness.

**Highlights:**

The first reported attempt of the Dream Catcher test.
The correlates of conscious experience may not lie in EEG spectral power.
Reported markers of NREM dreaming consciousness misperformed in a blinded setting.
Those markers also could not be confirmed in an unblinded setting.

## Introduction

### Background

If we take consciousness to be a natural, biological phenomenon that depends on neural activities inside the brain, then there must be objective patterns of brain activity that directly constitute consciousness and therefore correspond to the subjective features of experience. Whatever type of neural activity consciousness turns out to be, it must be something that incarnates the spatiotemporal patterns of phenomenal experience.

The Dream Catcher test was introduced as an empirical criterion for what would constitute a genuine scientific discovery of the underlying neural constituents of phenomenal consciousness (Revonsuo, 2006). The present empirical study, which we call the Dream Catcher *experiment*, is the first attempt to execute such a test. The test itself was originally an idealized thought experiment, devised to address the explanatory gap that exists between the physical explanation of consciousness and the phenomenal experience of consciousness itself—the so-called “hard problem” (Chalmers, 1995; Levine, 1983). Arguably, even if we were able to identify the neural correlates of consciousness, these would not suffice to bridge the explanatory gap: *correlations* do not in themselves provide an *explanation* for a phenomenon. Revonsuo (2006) proposed that consciousness would be genuinely explained by the discovery of *constitutive* mechanisms of consciousness at the phenomenal level. For the crucial distinction between correlates and constituents, see also Revonsuo (2001) and Miller (ed.) (2015).

To determine the constitutive mechanisms of consciousness, the Dream Catcher test requires researchers to make predictions about the qualitative features of participants’ phenomenal experience through use of purely brain-based data, without access to any information about the participant’s stimulus environment, subjective experience, or correlated brain patterns to known perceptual stimuli. This is achieved by the following stipulations. First, the study of consciousness is restricted to the domain of sleep to ensure that the contents of consciousness are largely independent of external stimuli. Second, the researchers charged with testing their brain-based model of consciousness are blinded to participants’ subjective reports, which are instead recorded by an independent team. Together, these restrictions bring the design closer to a no-report paradigm (Tsuchiya, Wilke, Frässle, & Lamme, 2015), which aims to prevent conflation between processes underlying conscious experience and those underlying the act of reporting conscious experience.

Conscious mentation is not just frequent in REM sleep but also occurs throughout the majority of non–rapid eye movement (NREM) sleep (Nielsen, 2000; Nir & Tononi, 2010; Noreika, Valli, Lahtela, & Revonsuo, 2009; Windt, Nielsen, & Thompson, 2016). Recently, specific spectral changes in sleep electroencephalography (EEG) have been found in studies contrasting periods of NREM sleep associated with reports of dreaming against periods without dreaming (Chellappa, Frey, Knoblauch, & Cajochen, 2011; Esposito, Nielsen, & Paquette, 2004; Scarpelli et al., 2017; Siclari et al., 2017; Siclari, Bernardi, Cataldi, & Tononi, 2018; Siclari, LaRocque, Bernardi, Postle, & Tononi, 2014). (For a review, see Ezquerro-Nassar & Noreika, 2019.) These studies concurred that reduced low-frequency EEG power correlates with dream recall, although they neither agreed on the source location of this difference, nor on whether high-frequency activity was also correlated with dream reports. We decided to put these findings under stricter scrutiny in our study, implementing the first reported attempt of the Dream Catcher test.

The Dream Catcher test’s proposed blinding aspects elevate it above most conventional approaches to the study of consciousness. Blinding also distinguishes our study from the previous studies, cited earlier, on the neural correlates of dreaming. Finally, the disconnection between external stimuli and subjective experience during sleep further prevents the conflation between conscious experience and external stimulus processing, which—while central to waking perception—plays a minimal role in dreams.

For our study, the Dream Catcher experiment, we have highly simplified the original Dream Catcher test due to the current limitations of neuroscientific knowledge and brain activity measuring capability. Instead of requiring researchers to reconstruct the content of phenomenal experience from a comprehensive set of brain activity data, we introduced a more realistic requirement: to identify the presence vs. absence of dreaming (i.e., the presence vs. absence of conscious experience) from polysomnograms, without access to information on whether participants had reported a dream after awakening or not. Although some studies have investigated neural correlates of specific dream content (Dresler et al., 2012; Horikawa & Kamitani, 2017; Horikawa, Tamaki, Miyawaki, & Kamitani, 2013; Siclari et al., 2017), reconstruction of the full phenomenal level would be a step for the distant future.

### Measures of consciousness

Early polysomnography (PSG) measured voltage fluctuations at various sites of the body, traced onto continuous paper feed, to be interpreted and classified by researchers by eye. Based on such features as the frequency of oscillations at the scalp, intensity of muscle tone activity and type of eye movement, researchers found that they could classify distinct stages of sleep and correlate them with dream reports (Aserinsky & Kleitman, 1953; Jouvet, 1967). Researchers have since increased the array of tools for analysing these same data, including spectral methods, phase coherence measures and the vast variety of methods devoted to time series analysis in general (Arsiwalla & Verschure, 2018; Cohen, 2014). Many features of brain electrophysiology have been investigated and reported to correlate with different conscious processes or even the level of consciousness. Spectral power differences have been commonly found at characteristic frequency bands; notably, lack of consciousness has been associated with increased power at low frequencies (delta waves: <4 Hz) in multiple contexts, including sleep stage depth, dream recall within a sleep stage, and anaesthetic depth (Chellappa et al., 2011; Evans, 2003; Hobson & Pace-Schott, 2002; Murphy et al., 2011; Scarpelli et al., 2017; Siclari et al., 2017, 2018; Thomsen, Rosenfalck, & Nørregaard Christensen, 1991). Higher levels of consciousness (or arousal) have also been suggested to correlate with a lower spectral exponent (Colombo et al., 2019), higher signal entropy or complexity (Bein, 2006; Bruhn, Röpcke, & Hoeft, 2000; D’Andola et al., 2017; Hudetz, Liu, Pillay, Boly, & Tononi, 2016; King et al., 2013; Liang et al., 2013; Ouyang, Li, Liu, & Li, 2013; Sarasso et al., 2015; Schartner et al., 2015), stronger phase coherence between brain areas (Bola et al., 2017; Lee et al., 2017; Mikulan et al., 2017), and more causally integrated brain areas (Barrett et al., 2012; D’Andola et al., 2017; Fasoula, Attal, & Schwartz, 2013).

With an ever increasing number of methods, we must be wary that almost surely a proportion of reported effects will be false positives. Particularly in cognitive neuroscience and psychology, the high prevalence of unreplicable studies has been a serious issue (Fanelli, 2009; Kriegeskorte, Simmons, Bellgowan, & Baker, 2009; Schooler, 2014; Vul, Harris, Winkielman, & Pashler, 2009). In this regard, a virtue of the Dream Catcher test is its blinded nature; it prevents biasing researchers towards a certain outcome due to knowing the true conditions of their samples. Thus, while the Dream Catcher test can generally be considered a paradigm for confirming the constitutive mechanisms of phenomenal consciousness, in our experiment it can be considered to confirm reported measures of the presence vs. absence of consciousness.

### Study design

The Dream Catcher experiment involved two teams. The first team was composed of Valdas Noreika, Levente Móró, Antti Revonsuo, and Katja Valli, who designed and collected data for the overall experiment and the Dream Catcher protocol—we’ll call this the Data Team. The second team was composed of William Wong, Jennifer Windt, and Naotsugu Tsuchiya, who analysed and classified the brain-based data with restricted access to participants’ dream reports—we’ll call this the Analysis Team. At the beginning of the experiment the Analysis Team only knew (a) published details of the Dream Catcher’s data collection method using the early night serial awakening protocol (Noreika et al., 2009), (b) the scientific literature on dreaming and consciousness published at the time (pre-2018), (c) the instruction sheet (Supplementary Document 4), and (d) some additional background information from occasional email exchanges with the Data Team regarding the Dream Catcher procedure.

The Dream Catcher experiment is a first step towards carrying out the core idea of the Dream Catcher test by focussing on dreams during carefully matched NREM sleep stages 2–3. By contrasting the recorded brain activity between dreaming and non-dreaming states in NREM sleep, we expected to better isolate the effect of the presence vs. absence of dreams in the data. Unlike in REM sleep, which has a dream recall prevalence of about 80% (Hobson, Pace-Schott, & Stickgold, 2000; Nielsen, 2000), the frequency of dream reports obtained from stage 2 NREM sleep is roughly equal to that of non-dream reports (Nielsen, 2000; Noreika et al., 2009). NREM dreams tend to be more fragmented, thought-like, and less vivid than REM sleep dreams (Mutz & Javadi, 2017). There is a contention that the non-vivid sleep mentation in NREM sleep should be categorised separately and that only multimodal, narratively complex, and often emotional experiences, which are typical of REM sleep, should be classified as dreaming (Hobson et al., 2000). However, as we are interested in the presence vs. absence of even minimal forms of consciousness, we shall refer to all reports of mentation during NREM sleep as dreams in this paper (for the detailed discussion of this theoretical position, please see Discussion).

Please note the unusual structure of our paper, which stems from our complex experimental setup. In the *General Methods* section, we describe the Data Team’s data collection procedures and the Dream Catcher experiment protocol, and give an overview of the Analysis Team’s strategy for blind classification. The particular procedures and results of the Analysis Team at each blinded step are described in the section *Blind Classification Methods and Results*, which is written from the Analysis Team’s point of view. This is followed by the *Post Hoc Evaluation* section, which describes the post-experimental analyses we performed to give further context to our results. We close with a discussion of the theoretical and methodological implications of the results for the Dream Catcher paradigm and dream research.

## General Methods

Study design, data collection and the blinding procedure were performed exclusively by the Data Team before any contact with the Analysis Team. The study protocol was approved by the Ethical Board of the University of Turku, and all participants signed informed consent following the Declaration of Helsinki. Data collection was conducted at the Sleep Laboratory at the Centre for Cognitive Neuroscience at the University of Turku.

Dream reports and PSGs were collected following an early night serial awakening paradigm (Noreika et al., 2009). Refer to Supplementary Document 1 for our sleep data collection procedures and methods, Supplementary Document 2 for the interview procedure, and Supplementary Document 3 for transcribed, exemplar dream reports.

### Participants

Fifteen Finnish-speaking volunteers were recruited to the study. They were screened to have no issues with psychological and neurological health, take no psychoactive drugs or have any sleep disorders at the time of the study. Aiming to assess the participants’ sleep latency and ability to give clear dream reports as well as to familiarize them with sleep laboratory environment, participants spent one adaptation night in the sleep laboratory. Five participants were excluded following adaptation nights due to sleeping difficulties in the laboratory, unclear dream reports upon awakening from NREM sleep, and/or sleep EEG artefacts due to sweating. The remaining 10 participants spent 4 experimental nights in the laboratory, for which each participant was compensated in total with 100 euros.

One of these participants did not recall any dreams upon awakening from NREM sleep, and hence this person’s data were not used in the Dream Catcher experiment. Thus, the final data utilised in the study was collected from 9 participants (4 males), aged 21 to 34 years (*M* = 27, *SD* = 5.39). Handedness was tested by means of the Edinburgh Handedness Questionnaire (Oldfield, 1971): eight of the participants were fully right-handed and one was fully left-handed.

### Data selection and blinding

All collected dream reports were divided by two blind raters (Master students in psychology) into four categories: 1) dreamless sleep, 2) white dream, 3) uncertain, and 4) dreamful (following Dement, 1955). For the Dream Catcher experiment, only the 1) dreamless and 4) dreamful categories were considered. The lowest number of reports from either category from a single participant was 3. Following this constraint, 3 dreamful and 3 dreamless sleep reports were selected from the 9 participants, yielding a pool of 54 reports and corresponding 1-minute pre-awakening EEG segments. The following criteria were applied in the data selection: (a) Both blind raters should have independently agreed on the basic recall category of the report; (b) in order to reduce the variability of reports (which might be reflected in the underlying brain activity to be blindly classified by the Analysis Team), all included dreams should be static (i.e., Orlinsky’s Modified Scale for Perceptual Complexity of Dreams, categories 1–4, 89% belonged to categories 3 & 4; Orlinsky, 1962; Noreika et al., 2009): we aimed to generate a perceptually homogeneous sample of dream reports, whereas the proportion of dynamic dreams is generally very low during early night NREM sleep; (c) the 60-s pre-awakening EEG should contain only NREM Stage 2 and/or Stage 3 epochs (i.e., three consecutive such 20-s epochs); (d) there should be comparable within-participant and between-participant proportions of Stage 2 and Stage 3 pre-awakening epochs across recall categories; and (e) selected EEG recordings should have a minimal amount of artefacts.

The blind classification phase of the study was to be undertaken by the Analysis Team. Data blinding for the Analysis Team was performed using a custom Perl script. This 219-line code loaded the metadata (such as the original recording number, the original participant number, the Session number 1 to 4, and the Condition “dreamful” or “dreamless”) from a comma-separated values file describing the parameters of the 54 samples. The recordings were randomly assigned labels with a consecutive numerical range: a general recording label (ID01–ID54), a participant label (S1–S9), a dreamfulness condition label (C1–C2), a participant-grouped condition label (G01–G18; i.e., 9 Participants × 2 Conditions), and a pairing label (P01–P27; i.e., 9 Participants × 3 Sessions) for pairs of dreamful vs. dreamless recordings from the same participant under the same condition. All these labels were logged into a Microsoft Excel table to be used by the Data Team for evaluating the Analysis Team’s blinded results. Finally, the script output a Windows batch file that renamed the original recording files to their randomised ID01–ID54 file names, to be received by the Analysis Team.

## Blind Classification Methods and Results

We refer to the individual 1-minute PSG recordings, given to the Analysis Team, as “cases”. In Table 1, we present a review of what information was given to the Analysis Team at each step of this blind classification task, as well as the terms we use in this article to refer to the various groupings of the cases revealed during the experiment. The terms are also illustrated in Figure 1.

**Figure 1.**
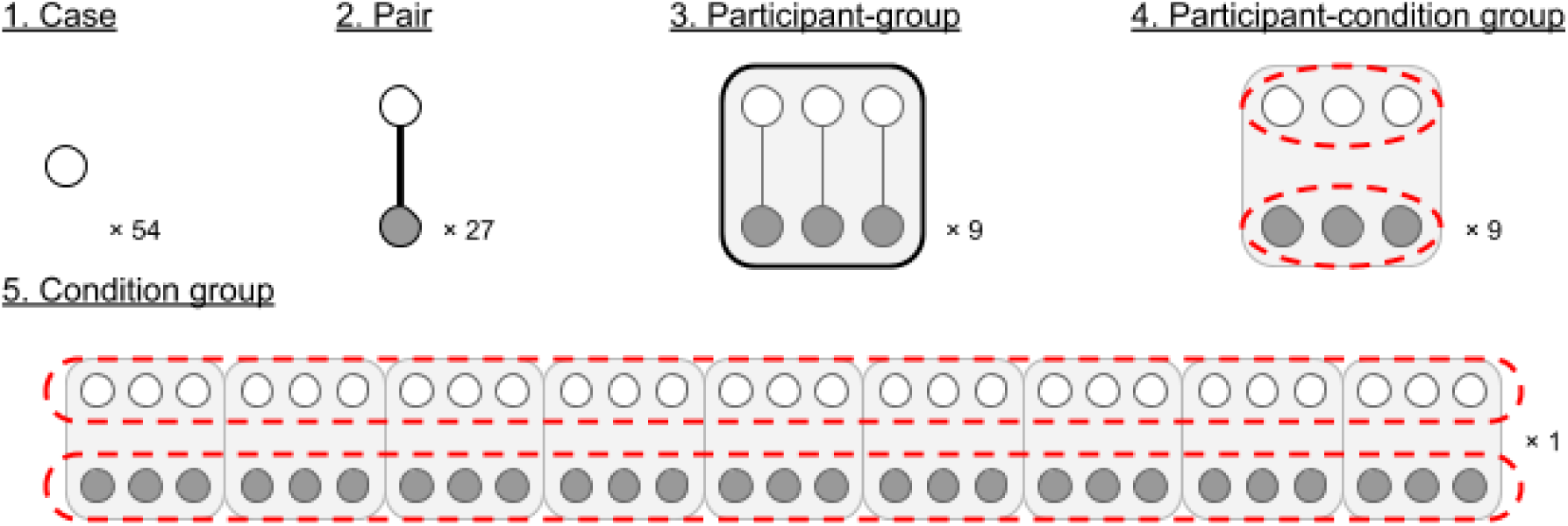
Illustration of blinding information at each step of classification.

**Table 1.** Blinding of information at each step of classification

| Step no. | Collective name for data | Available case information to Analysis Team |  |  |  |
| --- | --- | --- | --- | --- | --- |
| 1 | “Case” | - | - | - | - |
| 2 | “Pair” | Pairwise:<br>dreamful vs.<br>dreamless, same<br>participant. | - | - | - |
| 3 | “Participant-group” | Pairwise:<br>dreamful vs.<br>dreamless, same<br>participant. | Groups of 6<br>cases: same<br>participant. | - | - |
| 4 | “Participant-condition<br>group” | Pairwise:<br>dreamful vs.<br>dreamless, same<br>participant. | Groups of 6<br>cases: same<br>participant. | Groups of 3<br>cases: same<br>dreamfulness,<br>same participant. | - |
| 5 | “Condition group” | Pairwise:<br>dreamful vs.<br>dreamless, same<br>participant. | Groups of 6<br>cases: same<br>participant. | Groups of 3<br>cases: same<br>dreamfulness,<br>same participant. | All dreamful vs.<br>all dreamless. |

### Overall strategy

The methods described in this section were devised independently of the Data Team. The Analysis Team approached the classification problem firstly as a clustering problem. They assumed that brain states would be more homogeneous during non-dreaming than during dreaming, possibly because various contents of dreaming would diversify brain states. Such homogeneity would be amenable to cluster analysis, which is ideally suited to discover and group observations of high similarity in an objective manner, based on extracted features of the data. Only in the last step, where the two final clustered groups were classified for dreamfulness (a choice with only two alternatives), would the Analysis Team make a subjective determination of dreamfulness in line with previous findings.

The Analysis Team operated primarily in the Matlab software environment (The MathWorks, Inc., 2012); some of the EEG data handling was facilitated by the EEGLAB toolbox in Matlab (Delorme & Makeig, 2004). We will specify otherwise where relevant.

### Clustering Method

For clustering algorithms, the Analysis Team chose an evidence accumulation clustering (EAC) approach (Fred & Jain, 2005) with modifications. It was chosen over more common clustering techniques, such as *k*-means clustering and hierarchical clustering, for its demonstrated improved ability to identify clusters of arbitrary shapes and sizes. Throughout the blind classification procedure, the Analysis Team clustered cases into two groups based on the similarity of extracted features. For specific purposes of the study, they made changes to EAC. See Figure 2 for a comparison; details in Wong & Tsuchiya (in prep.). Here, we provide only a cursory description of the adopted EAC method.

**Figure 2.**
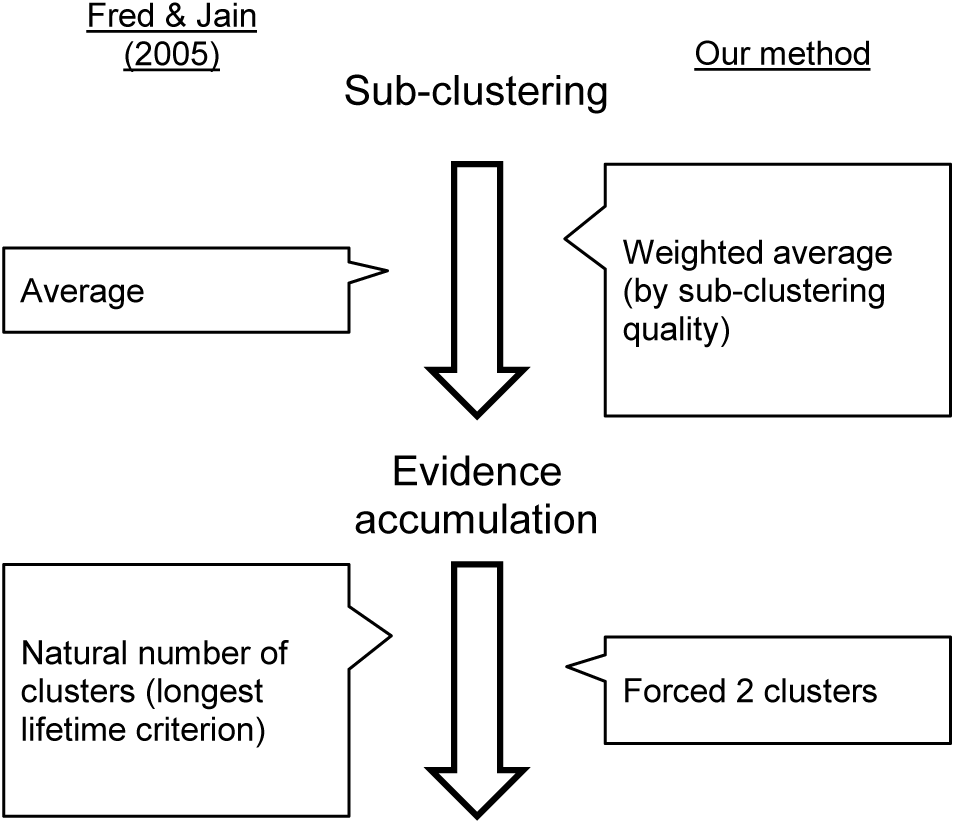
Contrast between Fred and Jain’s evidence accumulation clustering method and our method.

EAC works by accumulating the results of multiple clusterings—evidence—of the same data. Evidence consists of multiple runs of simpler clustering algorithms, where each may give incomplete but diversified information about similarities in the data. Here, we shall call these *sub-clusterings* so as not to confuse them with the hierarchical clustering procedure that follows the *evidence accumulation* step. In Fred and Jain’s experiments, the authors implemented sub-clustering using multiple, randomly seeded runs of *k*-means clustering (MacQueen, 1967) and then counted the frequency of co-association between each pair of data points. In the evidence accumulation step, these sub-clustering results were summed to produce the co-association similarity matrix, upon which they performed evidence accumulation via hierarchical clustering, with either single- or average-linkage criterion (Sokal & Michener, 1958). They finally thresholded the linkage distance so as to obtain the natural number of clusters present in the data.

The Analysis Team modified EAC for three reasons. First, unlike in the original EAC, the team knew from the outset that they were dealing with two equal-sized clusters. They therefore modified the final step from finding the natural number of clusters in the data to finding just two clusters. However, especially for noisy data sets, a linkage threshold that produces two clusters often produces ones of asymmetric sizes, wherein the smaller cluster may consist of one or a few outliers. To cope with this problem, the Analysis team chose the highest threshold that produced at least one cluster that is as close to 50% of the cases as possible, and all other clusters were reclassified as belonging to the other class. This forced the clustering results to produce two classes of close-to-equal sizes, where at least one of them was guaranteed to contain cases with similar features (i.e., are more homogeneous); the Analysis team postulated that this might represent dreamless cases.

Second, as the feature spaces consisted of up to 2,475 dimensions (or features), this far exceeded the maximum of 64 dimensions demonstrated by Fred and Jain. To avert a possible curse-of-dimensionality problem (see Bellman, 1957), the Analysis team limited the number of features to consider only up to nine at a time for each run. They populated the clustering ensemble with sub-clusterings of many different combinations of features.

Third, to give greater weight to more meaningful sub-clusterings during evidence accumulation, the Analysis Team also introduced weighting of sub-clustering results by a goodness-of-clustering metric.

The modified evidence accumulation clustering algorithm consisted of modifiable subcomponents: including sub-clustering, weighting, and hierarchical clustering. We will describe the specific methods used for each Step as they appear.

## Step 1

### Method

The task in the first step of blind classification was to classify 54 blinded 1-minute polysomnograms—referred to as *cases*—for dreamfulness. Data included simultaneous 25-channel EEG, 2-channel electrooculography (EOG) and 2-channel electromyography (EMG); the channels’ nominal locations were provided. The Analysis Team approached classification at this step with an exploration of feature extractions, followed by clustering based on a focussed set of features using the modified EAC (Wong & Tsuchiya, in prep.).

At this step, minimal information was known besides the data that could be used for classification. First, the Analysis Team considered eight different methods of analysis based on the previous EEG literature on levels of consciousness (e.g., sleep, anaesthesia, brain injury), listed with detailed methods in Supplementary Document 5. These were spectral power at established frequency bands, spectral power at fine frequency resolution, autocorrelation features as described by Thomsen et al. (1991), permutation entropy, approximate entropy, EOG root mean square (RMS) activity, EMG RMS activity, and spectral power in temporo-occipito-parietal areas (Siclari et al., 2014). As more features would be expected to result in overfitting (Domingos, 2012), the Analysis Team aimed to select only a few features fit for purpose.

The Analysis Team sought features that produced cluster results consistent with correct classification of dream report condition. In the absence of any ground truth, they looked into how consistently features clustered data when the data was split into four 15-s time segments. They looked for (a) clustering results that were consistent across temporally adjacent segments of time, or (b) results that were increasing in consistency for time segments more proximal to the time of awakening. A consistency metric between the clusterings of any two segments was formulated as follows.

Firstly, let us denote each case as *D*(*i*) (where *i* = 1, 2, …, 54), and divide it into 4 segments as *D_j_*(*i*) (where *j* = 1, 2, 3, 4). Clustering would assign to each *D_j_*(*i*) a membership label for one of two clusters: *c*1 and *c*2. Importantly, these clusters were not classified for dreamfulness; thus, *c*1 in one segment can correspond to either *c*1 or *c*2 in any other segment. But, because we always labelled the data by one of two clusters, there are only two possible ways to map *c*1 from one segment to *c*1 from another. Let us denote the proportion of cases that remain in the same cluster between segment *m* and *n* under the first and second mappings as *π*_1_(*m*,*n*) and *π*_2_(*m*,*n*). Then, the equality *π*_1_(*m*,*n*) + *π*_2_(*m*,*n*) = 1 holds for any clustering result. We thus define consistency of clustering between segment *m* and *n* as follows:

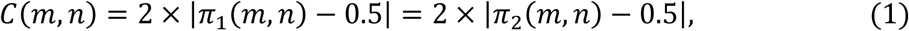

in which *C*(*m*,*n*) takes the value 0 when the clustering is not consistent at all, and 1 for perfect consistency.

The Analysis Team performed clustering on all cases for each time segment separately, and with each of the eight methods of feature extraction. Based on this consistency measure, in Step 1, they chose to classify dreamfulness based on the *PowerFine* feature set (detailed in Supplementary Document 5).

Briefly, the *PowerFine* feature set consisted of cases’ power spectral density (PSD) estimates for each EEG electrode, in frequency bins between 0 to 49.5 Hz in 0.5 Hz steps (i.e., 99 features per electrode). The feature set in total had 2,475 features for each case (99 x 25 electrodes). Note that PSDs were estimated throughout this study using fast Fourier transform and Welch’s method (Welch, 1967): Hann windows with 80% overlap. The Analysis Team performed the sub-clustering stage of EAC 82,475 times; 2,475 sub-clusterings were performed corresponding to each unique feature, and 10,000 random combinations of features were sub-clustered for each number of features *k* between 2 and 9 inclusively. Following the completion of the EAC procedure, the Analysis Team contrasted each feature between the two clusters by taking the Cohen’s *d* effect size (Cohen, 1988) of their values after log-transformation. Cohen’s *d* is calculated as 
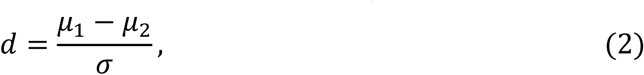
where the term *μ_1_* − *μ_2_* is the difference between the clusters’ means, and *σ* is their pooled standard deviation. The Analysis Team finally classified the cluster with overall higher low-frequency activity and lower high-frequency content to be from the dreamless condition, concordant with Siclari et al.’s (2014) findings.

### Results

The temporal consistency results are summarised in Figure 3*A*. The *PowerFine* candidate feature set exhibited highest consistency (> .9) for three pairs of consecutive time segments (i.e., 1 vs. 2, 2 vs. 3, and 3 vs. 4). This meant that class memberships of cases A and B tended to be consistent across four 15-s time segments, despite clustering being performed completely independently across segments. This and three other candidates were found significantly consistent following Bonferroni correction for multiple comparisons (two-tailed Binomial test, *N* = 54, unadjusted *p* ≤ .001), including *PermEn*, *Siclari* and *EmgRms*. With regard to positive trends in temporal consistency within individual participants, only *EmgRms* exhibited a difference between the temporal consistencies of the first half and last half (one-tailed permutation test, *p* = .0002, Bonferroni-corrected).

**Figure 3.**
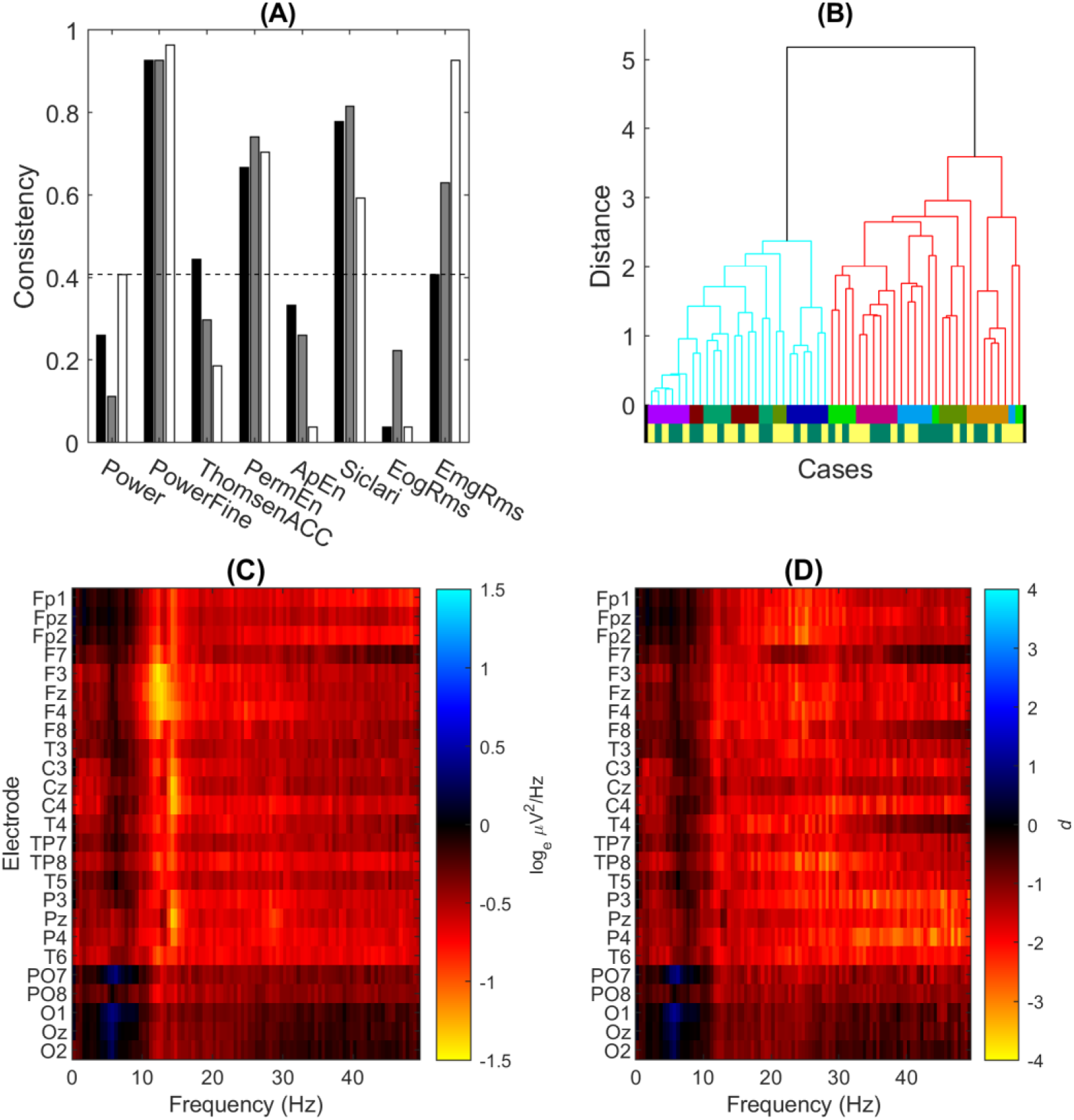
Step 1 blind classification results. (*A*) Temporal consistency for each of the 8 candidate feature sets. The consistency measure quantified the degree of agreement of unsupervised clustering between two consecutive 15-s segments of data; a consistency of 1 corresponded to identical clustering results. The black, grey, and white bars respectively correspond to the consistencies between segment 1 and 2, 2 and 3, and 3 and 4. The dashed line is the upper bound of the 95% CI for the null model, computed by Monte Carlo simulation, with Bonferroni correction. The Analysis Team decided to use the *PowerFine* feature set for Step 1 classification based on this result. (*B*) Dendrogram of clustering in the *PowerFine* feature set. The two clustered branches are differentially coloured bright cyan (Cluster 1) and darker red (Cluster 2). Cases were hierarchically clustered using UPGMA linkage, following evidence accumulation of their pairwise co-association similarity. The bottom two colour-coded rows are the true Participant identities (only revealed to the Analysis Team after Step 2) and Dreamfulness identities (only revealed after Step 5), respectively; cases with the Dreamful condition are in dark green, and Dreamless in bright yellow. (*C*) Mean difference in power spectra (Cluster 2 subtracted by Cluster 1) for all 25 EEG channels as a heat map. The colour scale is in units corresponding to the natural logarithm of μV_2_/Hz. (*D*) The effect sizes of the difference in *C*, quantified in Cohen’s *d*.

Using the *PowerFine* feature set, the Analysis Team performed modified EAC and obtained two clusters with unequal membership numbers (28 vs. 26). The dendrogram in Figure 3*B* gives a visualisation of the separation between the clusters. Figure 3*C* shows the mean difference in power by Cluster 1 subtracted from Cluster 2, and Figure 3*D* shows its effect size in Cohen’s *d*. The Analysis Team observed generally lower high-frequency power (ca. 11–30 Hz) in the average of Cluster 1 compared to Cluster 2 (visible in heat maps Fig. 3, *C*– *D*). Thus, based on Siclari et al. (2014)’s findings, the Analysis Team interpreted Cluster 1 to contain more dreamless report cases, and Cluster 2 to contain more dreamful ones. (To see the exact case-wise classification for this and all proceeding Steps, please refer to Fig. 9.) The Analysis Team submitted this classification, and the Data Team determined the accuracy to be 54%, which was not significantly different from chance according to a two-tailed Binomial test (*N* = 54, *p* = .68).

The Analysis Team’s clustering turned out to strongly group for participant identity and not for dreamfulness. Although full participant information was not revealed until Step 3, we show them in Figure 3*B* below the dendrogram by the upper row of coloured lines, whose colours code for each of the 9 participants. The fact that clustering more strongly grouped for participant identity was nonetheless determinable by the Analysis Team in Step 2: with the revelation of pairing information, it was found that 24 of the 27 pairs of cases were wrongly classified as co-occurring in the same cluster.

## Step 2

### Method

Together with the announcement of the result for Step 1, the Data Team removed the first layer of blindness by revealing their grouping into 27 *pairs* of cases. Each pair consisted of one dreamful and one dreamless case from the same participant. Still employing the modified EAC method, the Analysis Team was guided by a simple formalism of the linear mixed model:

Cases ∼ Dreamfulness + (1|Participant),

where the observed data (Cases) should reflect the main effect (Dreamfulness) with the added random effects (1|Participant). Assuming that the main effect of dreamfulness would be constant across participants, the Analysis Team treated the 54 cases as 27 single observations, each with feature values taken as the difference between a given pair. If these assumptions were correct, one can see that successful classification would be obtained through minimization of the variance across paired cases (i.e., alignment of feature vectors along the average difference between the two classes).

Also at this step, the Analysis Team reconsidered the features to use for clustering. In the previous step, their feature set consisted of 2,475 features of PSDs across the scalp, which they suspected in hindsight to have been excessively numerous and thus contributed to poor classification performance due to overfitting data (Domingos, 2012). They also suspected that the clustering algorithm might have performed better with a more encompassing feature set than one that only looked at EEG. To address these issues, the Analysis Team both condensed the EEG PSD features to a smaller number and expanded the diversity of features that composed the feature set for classification. (See Supplementary Document 6 for details.)

In brief, because the spectra across electrodes were apparently similar (see Fig. 3), the Analysis Team averaged the PSDs across all electrodes in 19 frequency bins that were logarithmically spaced—thus reducing the number of features from 2,475 to 19. As for the lost locality information, they delegated this to a focussed set of 11 features based on the hot zone findings reported by Siclari et al. (2014), which was accessible as a preprint manuscript at that point in time. These features included the whole-brain power at low and high frequency bands, low-frequency parieto-occipital power at various time windows, high-frequency frontal power at various time windows, and high-frequency power at hot zones relating to various perceptual categories. Lastly, the Analysis Team included 20 features extracted from the time course of EMG and EOG, computed differently from Step 1. For each 30-s segment of EMG or EOG, they computed the RMS values of all consecutive 1-s time windows, and took the 0^th^, 25^th^, 50^th^, 75^th^, and 100^th^ percentiles of those as features. This resulted in 10 features (5 percentiles × 2 segments) for each modality. In total, this set consisted of 50 features (19 scalp-average PSD + 11 *Siclari* features + 10 EMG + 10 EOG).

The difference between each pair was extracted by their subtraction in each feature value after Studentisation. Because the Analysis Team did not know which case of the pair belonged to which dream report condition, the polarity of the difference was arbitrarily assigned. As a result, they obtained 27 real-valued vectors in 50 dimensional feature space, whose *orientation*—and not direction—represented the difference between the pairs’ features.

Pairwise sub-clustering of the cases based on this transformed representation entailed the following. First, the feature space was subsampled to the combination of features to be used by the sub-clustering. Figure 4*A* shows a mock example where two features were chosen, thus representing the difference vectors (of which there are three, for illustrative purposes) in two dimensions. Next, each vector was centred at their midpoints and normalized to have unity length (Fig. 4*B*). A new vector representing the average orientation among all the difference vectors was then found (Fig. 4*C*), called the mean orientation vector, which should maximise the mean absolute cosine similarity between each difference vector and itself. Its orientation estimates the true qualitative difference between dreamfulness and dreamlessness. A hyperplane was then drawn normal to this vector, intersecting the origin (Fig. 4*D*). This hyperplane split each difference vector in two, which finally allowed those corresponding cases falling on each side of the hyperplane to be sub-clustered together.

**Figure 4.**
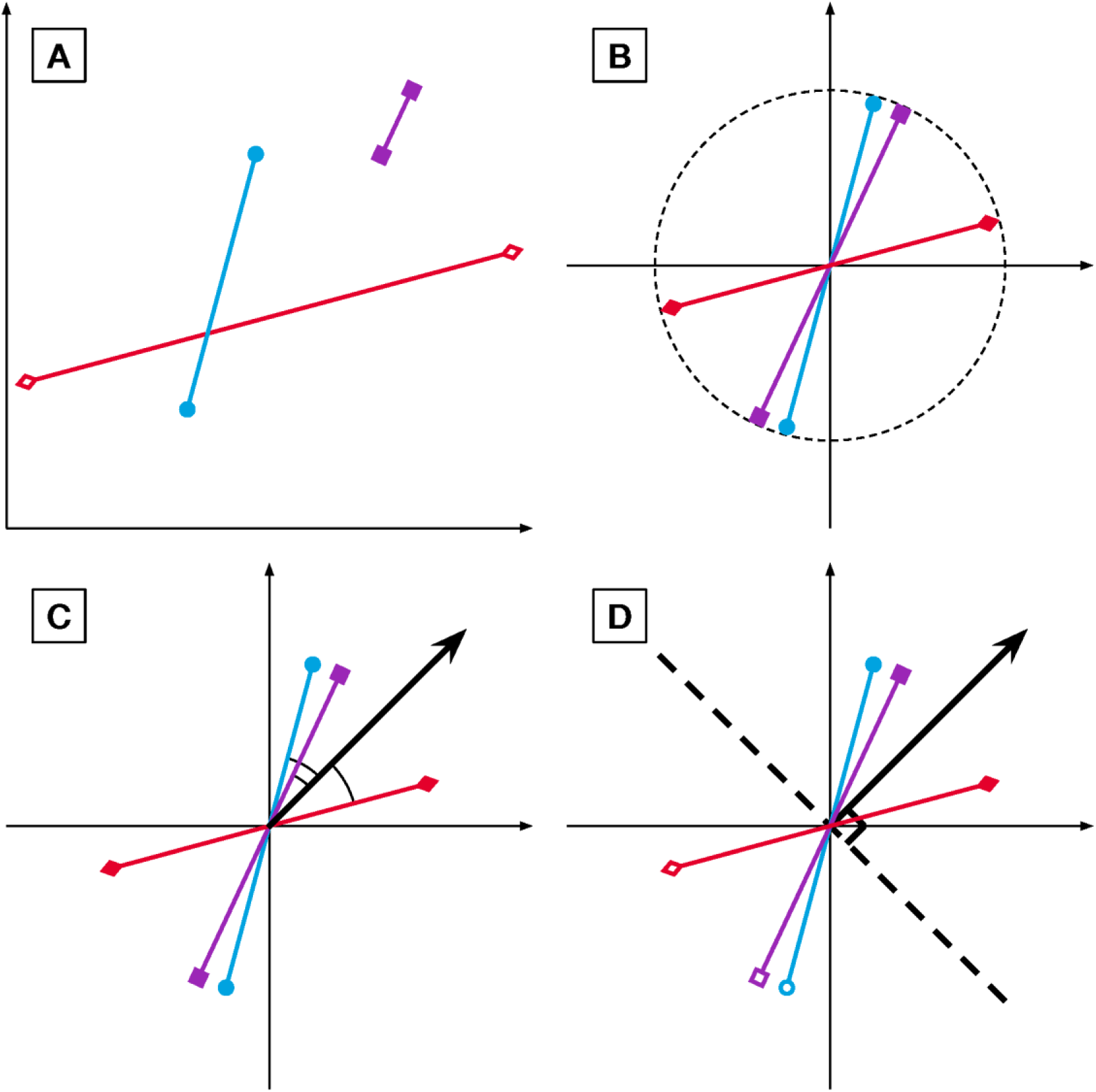
Step 2 pairwise sub-clustering schematic. Here, cases are depicted as points in a 2-D feature space; pairs are represented as two cases joined by a line. The basic concepts of pairwise sub-clustering begin in *A*, with the pairs of cases producing each a vector quantifying the difference between a dreamful case and a dreamless case. These difference vectors are then normalised in *B* to have equal length and intersect with the origin at their midpoints. They are now more representative of the qualitative difference between paired cases. Next, the mean orientation among all difference vectors is estimated in *C*, defined as a vector whose orientation maximises the average absolute cosine similarity between it and the difference vectors. The mean orientation vector estimates the true qualitative difference between dreamfulness and dreamlessness. It is depicted in the figure as a bold arrow originating from the origin. Finally, the cases are sub-clustered in *D* by introducing a hyperplane normal to this vector (bold, dashed line), and labelling the cases falling on each side of the hyperplane as co-associating. Paired cases will never co-associate.

For the goodness-of-clustering values, used to weight each sub-clustering within modified EAC, the Analysis Team took the mean absolute cosine similarity. In order to make these values comparable between sub-clusterings of different dimensionalities, the mean absolute cosine similarity was also divided by the expected value of such if the difference vectors were randomly oriented. This was computed using Monte Carlo simulations, where the number of difference vectors and the dimensionality were matched to those of each sub-clustering.

### Results

The Analysis Team obtained two equally sized clusters following the clustering procedure. The dendrogram in Figure 5*A* gives a visualisation of the separation between the clusters. Differences were found in the mean of their features most prominently in EOG activity (Fig. 5*C*); the cluster with higher EOG activity (Cluster 1) also had higher EMG activity (Fig. 5*B*) and low-frequency EEG activity (Fig. 5*D*). Differences in *Siclari* hot zone features were small and otherwise inconsistent (Fig. 5*E*). Although major differences were manifest in the EOG and EMG features, the Analysis Team had no rationale based on the literature for using them to determine which cluster corresponded to dreamfulness. Thus, based on EEG differences, they interpreted Cluster 1 to contain more dreamless report cases, and Cluster 2 to contain more dreamful report cases. The Data Team determined the accuracy of this classification to be 59%, which was not significantly different from chance (two-tailed Binomial test, *N* = 27, *p* = .44). Unlike after Step 1, the Analysis Team did not gain much new information from the feedback on their performance and the newly revealed participant identities.

**Figure 5.**
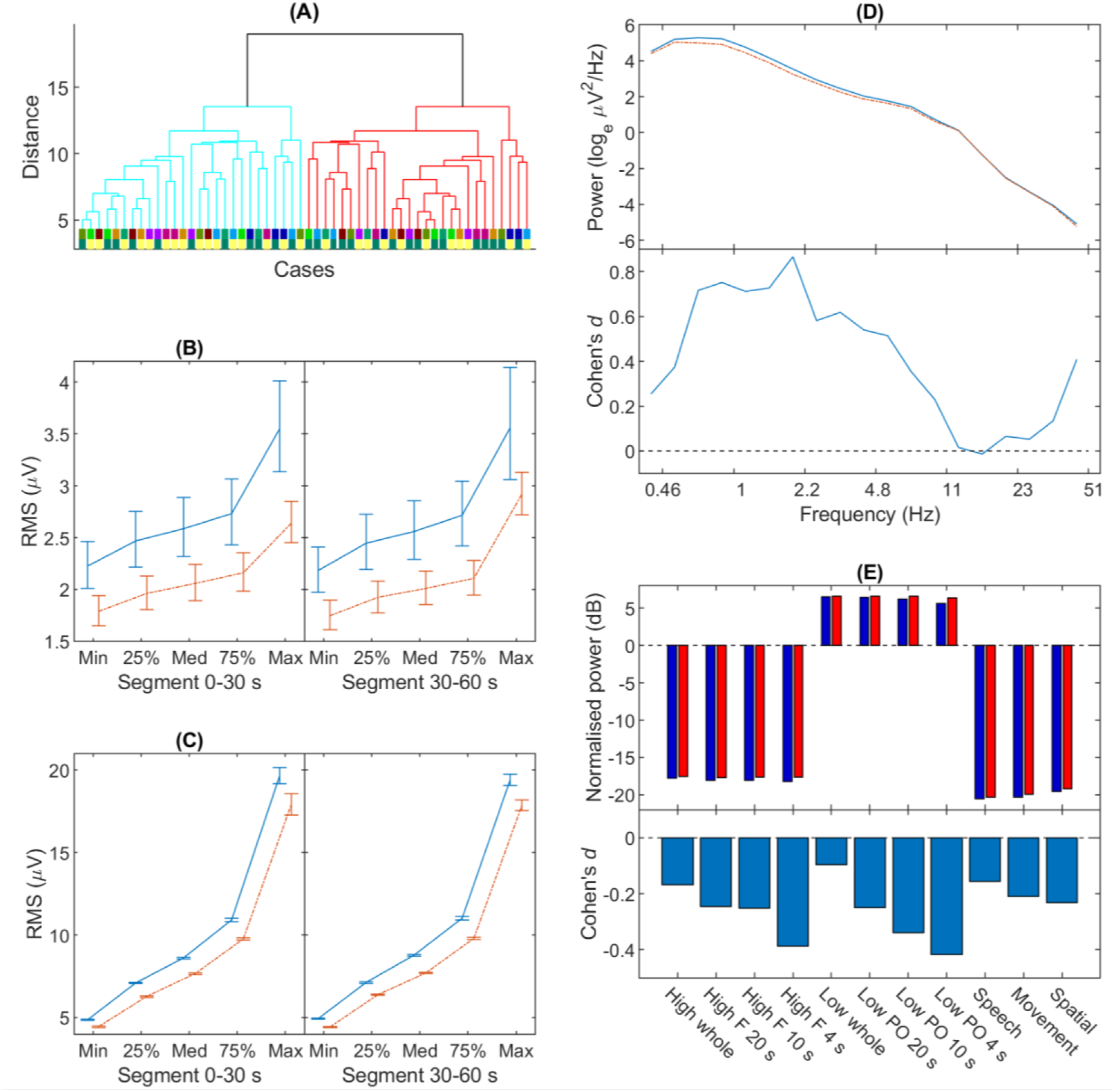
Step 2 blind classification results. *A*: Dendrogram of clustering, in the same format as in Figure 3A. Cases were hierarchically clustered using UPGMA linkage, following evidence accumulation of their pairwise co-association similarity. *B* shows the EMG feature means and standard errors of Cluster 1 (solid blue line) and Cluster 2 (broken red line) for the two time segments. Their statistics were calculated from log-transformed data. *C* shows the same for EOG activity. *D*: The EEG power spectral density average of Cluster 1 and Cluster 2, and their effect sizes. For the top panel, Cluster 1 is the solid blue line, and Cluster 2 is the broken red line. In the bottom panel, their statistical differences are expressed as Cohen’s *d*, in lieu of error bars for the top panel. *E*: The difference in power for frontal (“F”) and parieto-occipital (“PO”) electrodes, separately for “high” (18–50 Hz) and “low” (1–12 Hz) frequencies, over the indicated period of time just before awakening; and for the whole scalp locality (“whole”) over 20 s just before awakening. For the top panel, Cluster 1 in dark blue bars, and Cluster 2 in bright red. Like for *D*, their statistical differences are expressed in the bottom panel as Cohen’s *d*.

## Step 3

### Method

The Data Team removed the second level of blindness with the revelation of cases grouped by common participants. This resulted in nine distinct *participant-groups*, each associated with six cases, which were composed of three condition-balanced pairs of cases. Given this information, the Analysis Team exploited participant information by removing participant-specific, condition-irrelevant components from the EOG-EEG signals. To this end, they utilised independent component analysis (ICA), a technique for “unmixing” multichannel time series into their underlying, statistically independent time series components. They afterwards recomposed the case recordings after selecting and removing the condition-irrelevant components, which might have consisted of artefacts caused by eye and muscle movements. The full details of the methodology are described in Supplementary Document 6.

From the recomposed cases of all participants, the Analysis Team extracted the same set of features explained in Step 2 and performed EAC on it. A different set of EAC sub-clustering procedures was devised to take into account participant information. Taking the method of Step 2’s pair difference vector sub-clustering by mean orientation, the team sub-clustered the cases in four ways. In two of the ways, similar to Step 2, they sub-clustered cases in a pairwise manner: firstly with respect to the mean orientation amongst all pairs, and secondly with respect to the mean orientation of their own participant. The other two ways used the method of Step 1’s *k*-means sub-clustering and sub-clustered the unpaired cases: firstly amongst all cases, and secondly amongst each participant-group of cases. Therefore, there were four ways in which they performed sub-clustering. The Supplementary Figure S6.2 gives an overview of the scheme. They calculated the final similarity matrix as the average of these four different sub-clustering methods’ results; the exact details are also described in Supplementary Document 6. The resulting clusters produced with this method were thereafter classified for dreamfulness using the same rationale as in Step 1.

### Results

The Analysis Team obtained two equally sized clusters following the clustering procedure, using Ward’s method (Ward, 1963) as an alternate hierarchical linkage measurement. The UPGMA linkage method as in the previous Steps resulted in clusters of uneven sizes, which was undesirable. Figure 6 shows results for this step in the same format as Figure 5 for Step 2. The Analysis Team found differences between the clusters most prominently in EOG activity (Fig. 6*C*); the cluster with higher EOG activity (Cluster 2) also had higher EMG activity (Fig. 6*B*) and low-frequency EEG activity (Fig. 6*D*). In contrast, differences in *Siclari* hot zone features were small (absolute effect size Cohen’s *d* < 0.7) and otherwise inconsistent with each other in regard to their reported interpretations (Fig. 6*E*). Faced with similar results to Step 2, the Analysis Team interpreted Cluster 1 to contain more dreamful report cases (see Fig. 9 for exact case-wise classification).

**Figure 6.**
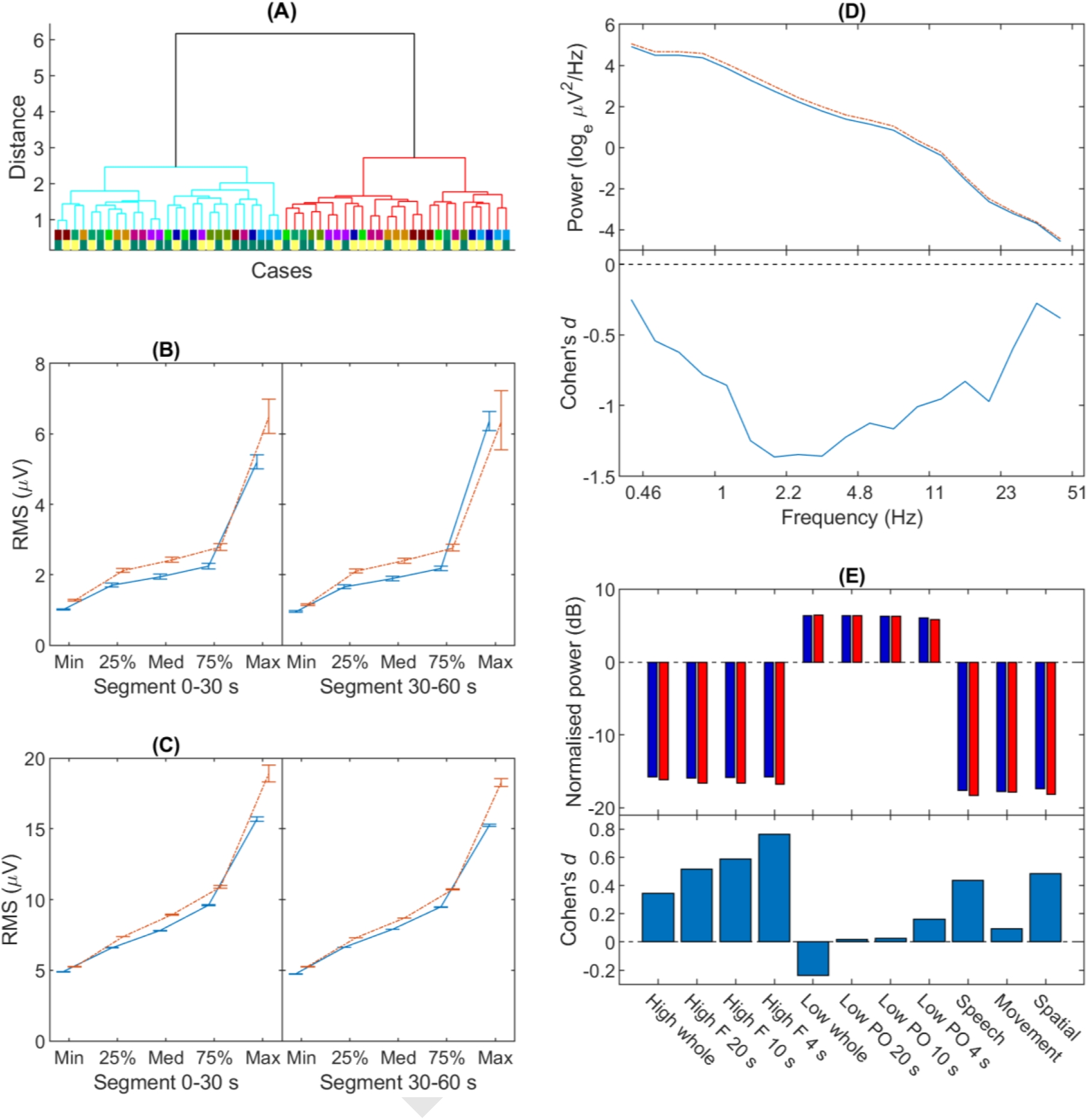
Step 3 blind classification results. The same format as in Figure 5. Cases were hierarchically clustered using Ward’s minimum variance linkage after evidence accumulation.

The Data Team determined the accuracy of this classification to be 59%, which was not significantly different from chance (two-tailed Binomial test, *N* = 27, *p* = .44). This feedback also did not give the Analysis team any further insights into their classification.

## Step 4

### Method

The third level of blindness was removed by grouping all cases with the same dream report condition from each participant. This effectively gave each participant two unlabelled condition groups of three cases each—referred to as *participant-condition groups*. The Analysis Team approached classification similarly to Step 3, first removing condition-irrelevant components using ICA with an adjusted component removal procedure, but then taking the difference in features between the average of conditions for each participant as observation vectors. This resulted in nine 50-dimensional vectors on which sub-clustering was performed in a pairwise manner. See Supplementary Document 6 for full details. Following EAC, the team classified the clustering result for dream report conditions according to the rationale set out in Step 1.

### Results

Figure 7 reports the results in the same format as in Figures 5 and 6. However, Figure 7*A* shows the dendrogram of clustering at only the participant level, unlike in Figures 5*A* and 6*A*, as they were already grouped at this Step. Unlike Steps 2 and 3, the Analysis Team found a prominent difference between the clusters in EMG activity (Fig. 7*B*) but not EOG activity (Fig. 7*C*); the cluster with higher EMG activity (Cluster 1) also had higher low-frequency EEG activity (Fig. 7*D*). Differences in *Siclari* hot zone features also indicated lower frontal high-frequency activity for this cluster (Fig. 7*E*), which in their study indicated an absence of dreaming experience. The team thus interpreted Cluster 1 to contain more dreamless report cases and Cluster 2 to contain more dreamful report cases (see Fig. 9 for exact case-wise classification). The Data Team determined the accuracy of this classification to be 44%, which was not significantly different from chance performance (two-tailed Binomial test, *N* = 9, *p* = 1).

**Figure 7.**
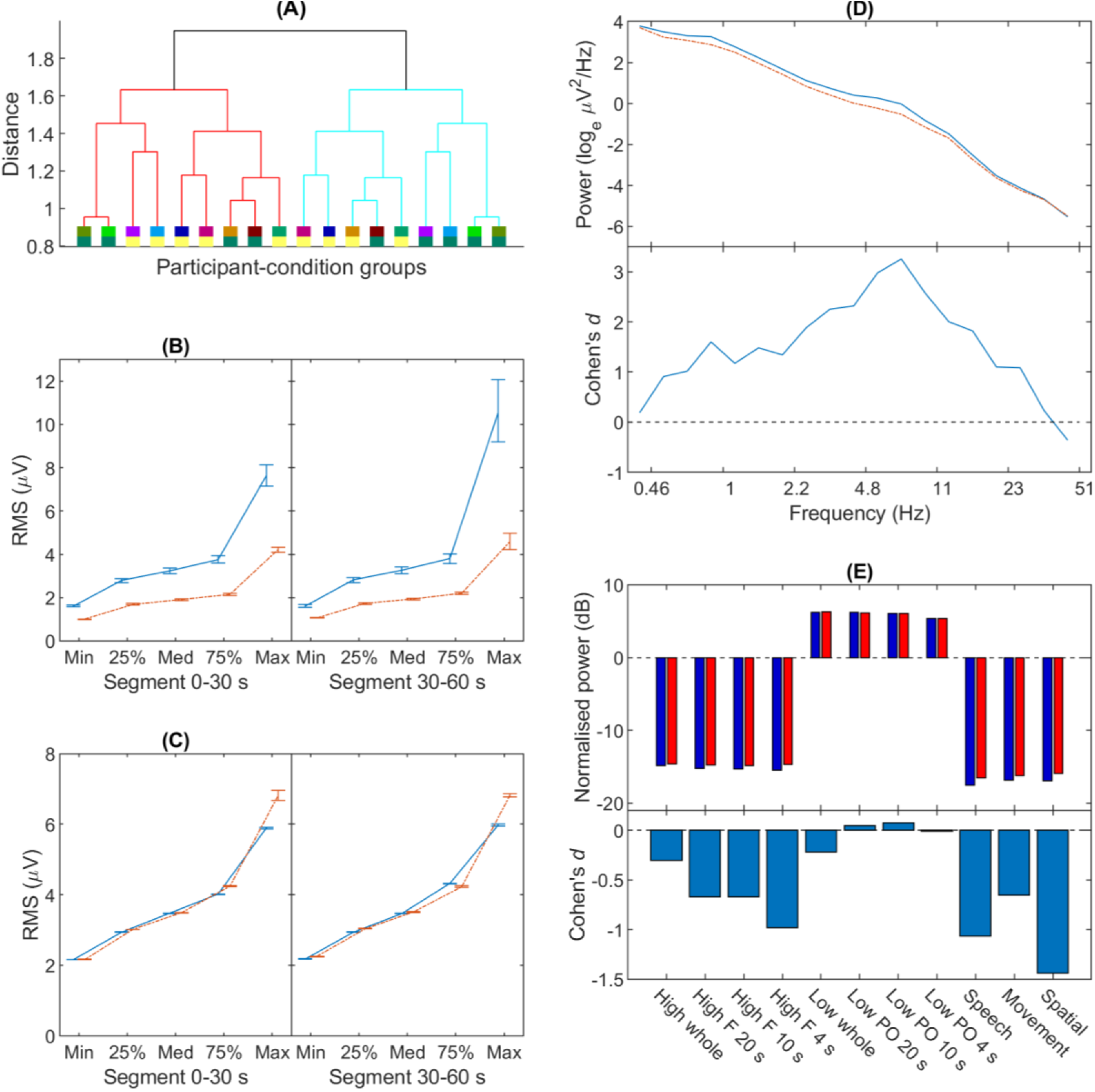
Step 4 blind classification results. The same format as in Figure 5 besides *A*, which shows the dendrogram on participant-condition group averages.

## Step 5

### Method

The penultimate layer of blindness to be removed was information on the common dream report condition across all cases. This resulted in just two groups—one from the dreamful condition, the other from the dreamless condition—each consisting of 27 cases fully labelled by common participants. No clustering was required. To make their final blind classification, the Analysis Team replicated three measures based on the significant differences reported by Siclari et al. (2017) and Scarpelli et al. (2017), both of which had been recently published. Two of the features were Siclari et al.’s low and high frequency hot zone power, named respectively *SBP low* and *SBP high* (SBP for the initials of the first three authors of the Siclari paper), and one feature was the low frequency activity reported by Scarpelli et al., named *Scarpelli* (see Supplementary Document 6 for full details). The Analysis Team classified these condition groups for dream report condition based on the group-average of these features.

### Results

The difference in feature means after removing inter-participant variability is shown in Figure 8*A*. All features had low effect sizes (absolute Cohen’s *d* < 0.4). As they were consistent in their indication of dream report condition, the Analysis Team interpreted Cluster 1 to contain more dreamless report cases, and Cluster 2 to contain more dreamful report cases (see Fig. 9 for exact case-wise classification). The Data Team determined this classification to be inaccurate.

**Figure 8.**
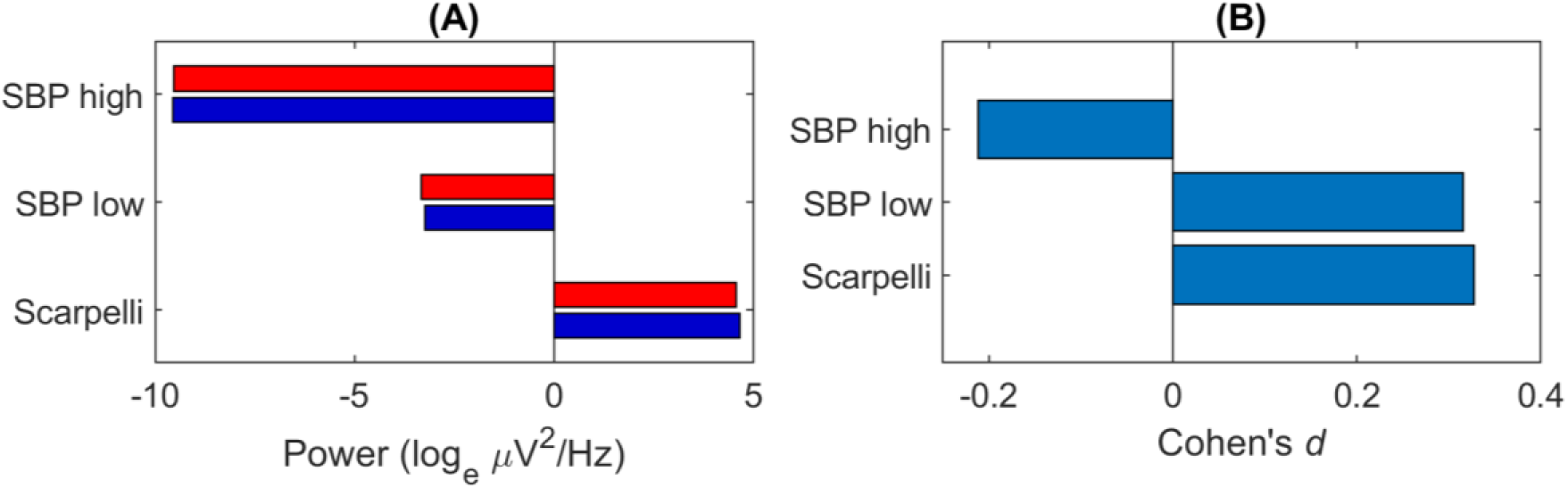
Step 5 blind classification results. (*A*) The difference in feature means between Cluster 1 and Cluster 2. Cluster 1 in dark blue bars, and Cluster 2 in bright red. (*B*) The effect sizes of the difference in *A*, of Cluster 1 from Cluster 2 in Cohen’s *d*, after subtracting each participant’s means.

**Figure 9.**
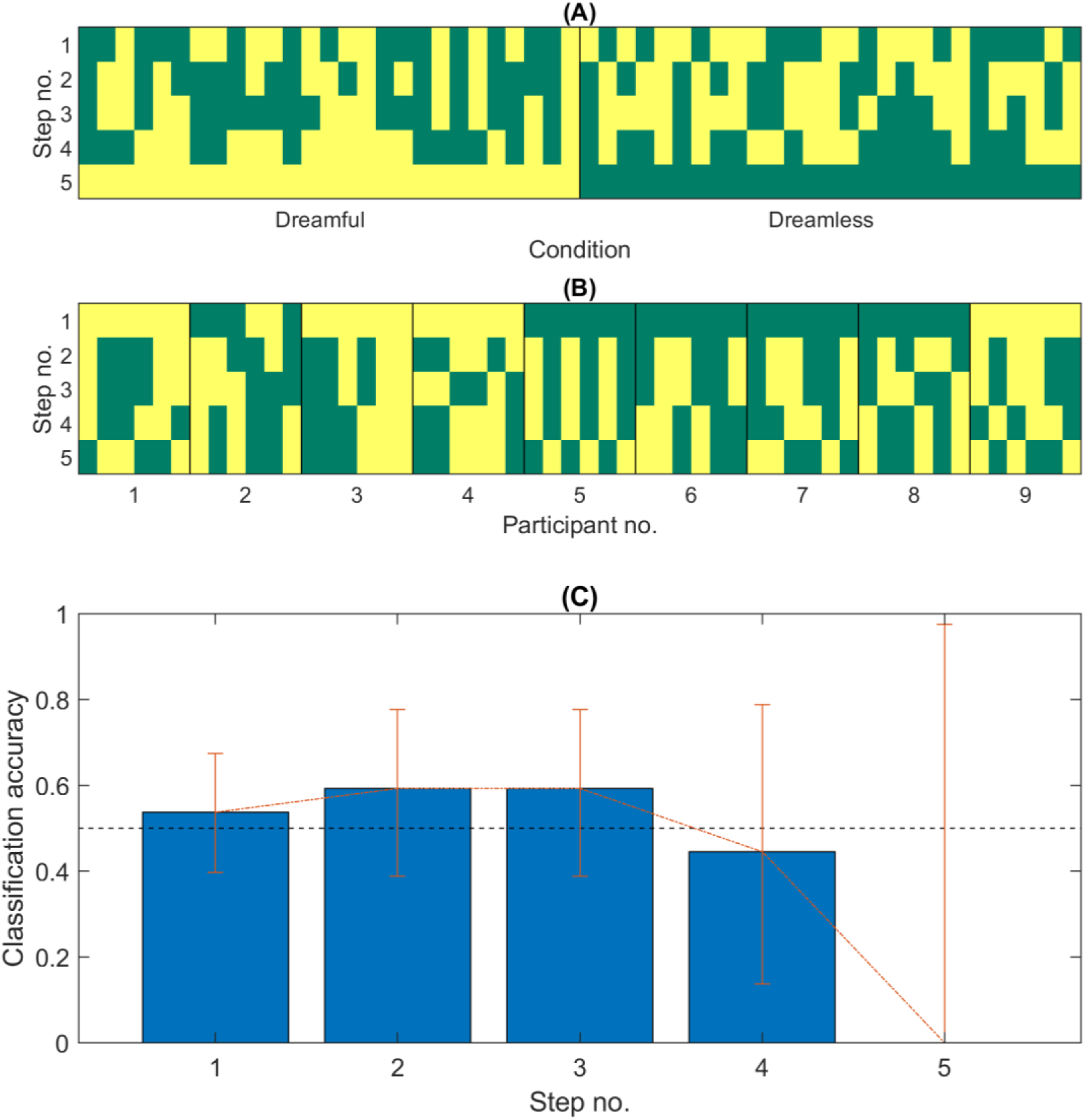
Case classifications through five levels of information. In *A* and *B*, the dark green cells are cases that were classified *dreamful*. Cases are horizontally sorted by Condition in *A* and Participant Number in *B*. In *C*, the error bars are 95% confidence intervals; chance-level accuracy (50%) is demarcated by the black, dashed line.

## Post Hoc Evaluation

From Steps 1 to 5, the Analysis Team never succeeded in classifying dreamful from dreamless sleep beyond chance-level performance (Fig. 9*C*). In the following post hoc analyses, we used all the information about the data and tried to quantify the effects of dreamfulness using more conventional, unblinded analysis methods. Through these analyses, we hoped to understand the source of the poor performance in the blinded classifications.

We set out to firstly quantify how well we could replicate selected effects reported in past studies (i.e. Scarpelli et al., 2017; Siclari et al., 2017) using Bayesian model selection. We also set out to quantify any other differences between the conditions via two methods: a multivariate analysis of variance (MANOVA) and a decoding analysis.

### Bayesian Model Selection

We investigated how our data compared with the then-recently reported effects of NREM dreaming, specifically in two 2017 articles by Siclari et al. and Scarpelli et al. We modelled the distribution of effect sizes expected for data of our sample size based on the reported effects using a Monte Carlo approach, and tested their plausibility in light of the replication attempts on our data against a null model with the use of Bayesian model selection (Goodman, 1999).

### Method

We reused features *SBP low*, *SBP high*, and *Scarpelli* from Step 5 of blind classification for replicating the effects being considered (see Supplementary Document 6 for full details).

We modelled past findings as probability density distributions of their effect sizes (Cohen’s (a) *d*) for data of our sample size *n* = 9 participants, with 3 dreamful and 3 dreamless data sets each. For each reported effect, we deduced the effect size based on the reported sample size and the *t*- or *F*-statistic. Then, assuming a Gaussian generator with this effect size, we stochastically simulated 10,000 experiments of *n* = 9 and obtained their resulting Cohen’s *d*s. We modelled the probability density distribution of *d* using Matlab’s *fitdist* function to obtain a Gaussian-kernel-smoothed density estimation, with an automatically chosen kernel width optimised for Gaussian densities.

Statistics for Siclari et al.’s (2017) posterior hot zone NREM findings for low and high frequencies were obtained from their Supplementary Figure 5: respectively *n* = 32, *t* = −2.98; and *n* = 32, *t* = 3.46. From this, their deduced Cohen’s *d*s were respectively −0.527 and 0.612. For Scarpelli et al.’s (2017) findings, we selected results from the nine electrode locations, reported for NREM recall vs. non-recall differences, that were statistically significant. We obtained the statistics from their Figure 1: *n* = 14, *F* ∈ (14.78, 15.49, 15.08, 15.94, 13.32, 19.45, 13.26, 13.35, 15.46). The deduced Cohen’s *d*s were (−1.03, −1.05, −1.04, −1.07, −0.975, −1.18, −0.973, −0.977, −1.05), and we took their average to get a single statistic *d* = −1.04.

In addition to Bayesian model selection, we also estimated traditional confidence intervals of the effect sizes of our data via bootstrapping of underlying features—constrained for the original numbers of participants and conditions—over 1,000 permutations.

### Results

The following results summarise those from Table 2. With respect to Siclari’s low frequency effect, our data’s effect size *d* = −0.39 supported this finding over the null model to a moderate degree (Bayes factor *K* = 3.3, 2 ln *K* = 2.4). However, in regard to Siclari’s high frequency effect, we found *d* = 0.29 for our data, meaning the null model was much more probable (Bayes factor *K* = 77^-1^, 2 ln *K* = −8.7). For Scarpelli’s low frequency effect, we found *d* = −0.56 for our data, which did not particularly support their finding but rather slightly favoured the null model (Bayes factor *K* = 2.6^-1^, 2 ln *K* = −1.9).

**Table 2.** Relative likelihood of Dream Catcher effect size given reported effect size

| Feature | $d$ | $d$ (95% CI) | $P(d\vert M_{Null})$ | $P(d\vert M_{Reported})$ | Bayes factor | |
| --- | --- | --- | --- | --- | --- | --- |
| | | | | | $K$ | $2 \ln K$ |
| Siclari low frequency | -0.39 | [-0.80, 0.19] | 0.31 | 1.0 | 3.3 | 2.4 |
| Siclari high frequency | 0.29 | [-0.34, 0.78] | 0.59 | $7.6 \times 10^{-3}$ | $1/77$ | -8.7 |
| Scarpelli low frequency | -0.56 | [-1.02, 0.26] | 0.31 | 0.12 | $1/2.6$ | -1.9 |

## MANOVA

We performed a MANOVA to parametrically discern whether there was a significant effect of dream report conditions amongst the features that we used during the course of blind classification.

### Method

We used a two-way MANOVA design that tested the effects (independent variables) of dream report condition, participant grouping, and their interaction. We combined all features used throughout our blind classifications and took these as our multivariate dependent variables. These were reduced from 2,628 to 18 dependent variables in order to make the MANOVA feasible, by limiting the degrees of freedom in the analysis. The number of reduced variables was chosen to minimise the Bayesian information criterion (Schwarz, 1978). For the reduction method, we performed singular-value decomposition on the features and applied a varimax rotation to the components; we took these steps to discourage variables from being composed of homogeneously mixed features and make the variables more independent of each other (Kaiser, 1958).

### Results

We tabulated the results of the MANOVA in Table 3. The sought-after main effect of the dream report condition on our features was not present (Lawley-Hotelling trace, *T* = 251, *p* = .93); however, we found a significant participant effect (Lawley-Hotelling trace, *T* = 2.9 × 10^4^, *p* < .001). We found no significant interaction effect between dream report condition and participants (Lawley-Hotelling trace, T = 232, *p* = .89).

**Table 3.** Two-way MANOVA of independent variables, participant grouping and dreamfulness grouping, on 18 SVD-reduced dependent variables of all extracted features

| Effect | $df$ (full) | $df$ (reduced) | Lawley-Hotelling trace | $p$ |
| --- | --- | --- | --- | --- |
| Dreamfulness | 36 | 45 | 251 | .930 |
| Participant | 36 | 52 | $2.86 \times 10^4$ | < .001 |
| Interaction | 36 | 44 | 232 | .889 |

### Decodability

To complement the parametric statistical analysis of MANOVA, we also performed a decoding analysis to discover the decodability of EEG power spectra using a support vector machine algorithm (Cortes & Vapnik, 1995).

### Method

We employed the nu-SVC model for our decoder, as implemented in LIBSVM for Matlab (version 3.18; Chang & Lin, 2011, 2014); its regularisation parameter was set to the default *nu* = 0.5, and the model was set to output probability estimates of classification. We validated this decoder as described in Supplementary Document 7. We estimated classification accuracy using participant-wise leave-one-out cross-validation on the data. For the 9 participants, each of whom acted as a fold for cross-validation, we classified each pair dichotomously—as we did in the blind classification experiments after Step 1—based on their relative probability estimate outputs. We preprocessed the PSD features by taking their natural logarithms, standardising them to have zero mean at each frequency bin in each participant and unit variance amongst all frequencies and participants, and then whitening the data using the zero-phase component analysis method for clustering using the Mahalanobis distance (Bell & Sejnowski, 1997; Mahalanobis, 1936).

First, we attempted to find the optimal time window length for decodability, since it was possible that our utilisation of the whole 60-s period in blind classification was suboptimal: in related studies, Siclari et al. (2017) used 20-s periods, and Nieminen et al. (2016) used 30-s periods; on the other hand, Esposito et al. (2004) used 3-minute periods, and Scarpelli et al. (2017) used 5-minute periods. We investigated this aspect by obtaining decoding accuracies of window lengths varying from 2 to 60 s. We used feature sets of common average electrode–referenced EEG PSDs for each of the 25 electrodes. PSDs were calculated using fast Fourier transform without zero padding, so that the number of frequency bins—and hence features—naturally scaled with window length.

Second, we searched for the optimal decoding performance of the same data amongst frequency bands and electrodes, as a function of time until awakening with a fixed window length. We chose to use 19 logarithmically spaced frequency bins, as in Step 2 of blind classification, and a window length of 4 s. PSDs were estimated using the multitaper method with 3 Slepian tapers to better capture the spectra across each whole time window, as they were non-overlapping. The spectral power of each desired logarithmic frequency bin was then taken to be the average power within that band. We then decoded the dream report condition at each frequency bin and time amongst the 25 electrodes and also at each electrode and time amongst the 19 frequency bins. We then visualised the results in a time-frequency and time-electrode heat map (Fig. 10).

**Figure 10.**
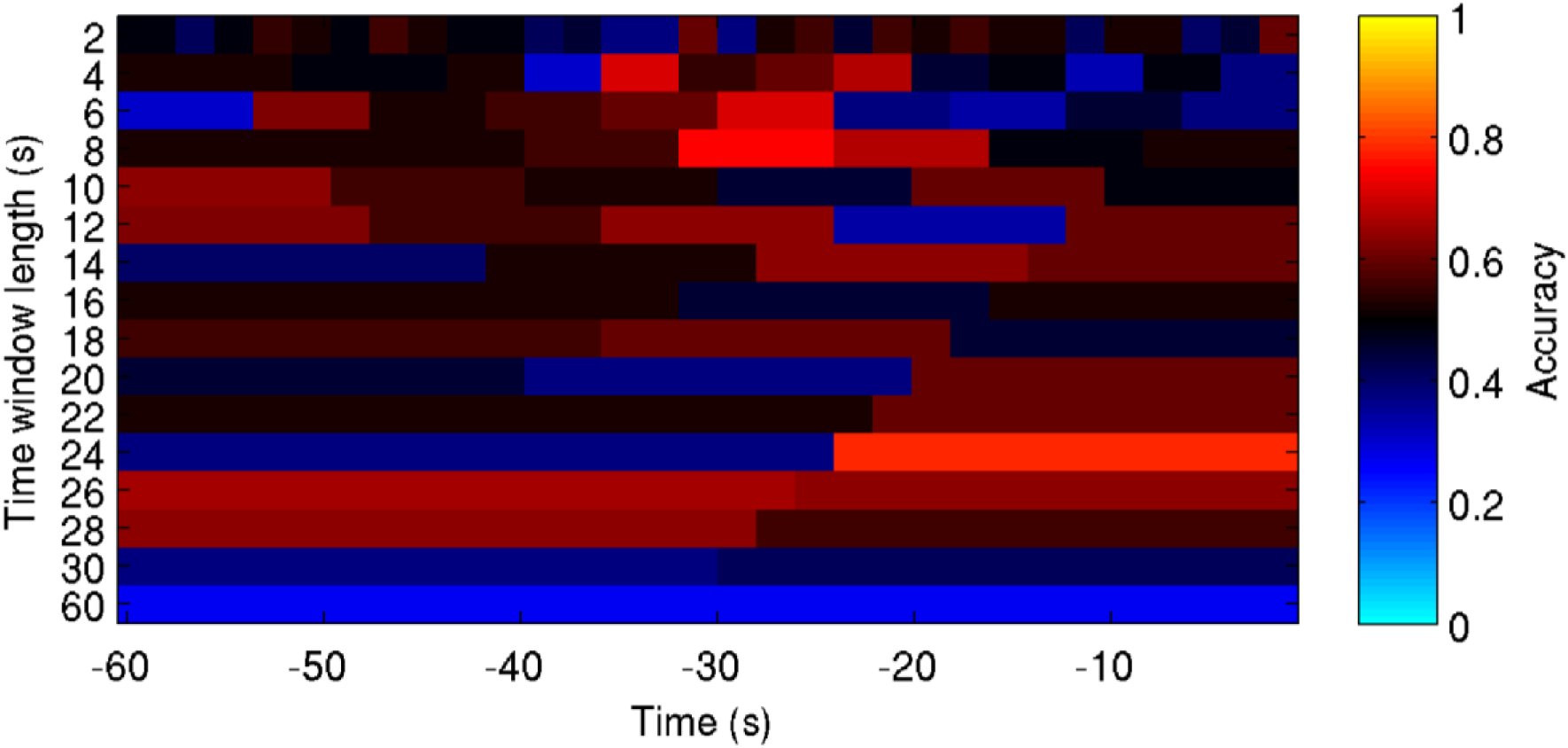
Dreamfulness decodability of electrode power spectral density features over various time window lengths and times. Accuracy of decoding is colour-coded, with chance-level performance corresponding to 0.5 accuracy (black). The critical thresholds for statistical significance at uncorrected *p* ≤ .01, .001 and .0001 respectively are accuracies .70, .78 and .85 (one-tailed Binomial test, *N* = 27).

We expected to find some combination of window length, time centre, electrode, and/or frequency band that would decode dreamfulness better than chance, assuming that a reliable difference is present in the data. We also expected decodability to increase for windows with time centres closer to awakening; such a result was suggested in Siclari et al.’s (2014) preprint manuscript.

### Results

The result of varying time window length on the decodability is shown in Figure 10. In general, decodability was not better than chance amongst all tested window lengths: (a) after adjusting for a false discovery rate *α* = .05 (Benjamini & Yekutieli, 2001), no average decoding performance within each test length was significantly better than chance (one-tailed Binomial tests, *N* = [27, 810], adjusted *p* ≥ 0.54); and (b) amongst all time windows contiguous with time of awakening, only one length of window (24-s) was significantly better than chance at 78% accuracy (one-tailed binomial test, *N* = 27, adjusted *p* = 0.04); furthermore, (c) no positive linear trend with time till awakening was apparent for any applicable length of window (one-tailed Pearson linear correlation, adjusted *p* > 1).

The time courses of decodability for 4-s windows with frequency- and electrode-wise partitioning are shown in Figure 11. Again, decodability was generally no better than chance: (a) one time window at 18 s prior to awakening had an average frequency-partitioned decoding performance that was significantly better than chance after adjusting for false discovery rate *α* = .05 (one-tailed Binomial test, *N* = 513, adjusted *p* = 0.04) at 57% accuracy, however this result was not corroborated by the variable window length results; (b) no frequency band had an average frequency-partitioned decoding performance that was significantly better than chance (one-tailed Binomial test, *N* = 405, adjusted *p* ≥ 0.94); (c) no positive linear trend with time was apparent for any frequency band (one-tailed Pearson linear correlation, adjusted *p* > 1); (d) no time window had an average electrode-partitioned decoding performance that was significantly better than chance (one-tailed Binomial test, *N* = 675, adjusted *p* ≥ 0.11); (e) no frequency band had an average electrode-partitioned decoding performance that was significantly better than chance (one-tailed Binomial test, *N* = 405, adjusted *p* ≥ 0.40); (f) no positive linear trend with time was apparent for any electrode (one-tailed Pearson linear correlation, adjusted *p* > 1).

**Figure 11.**
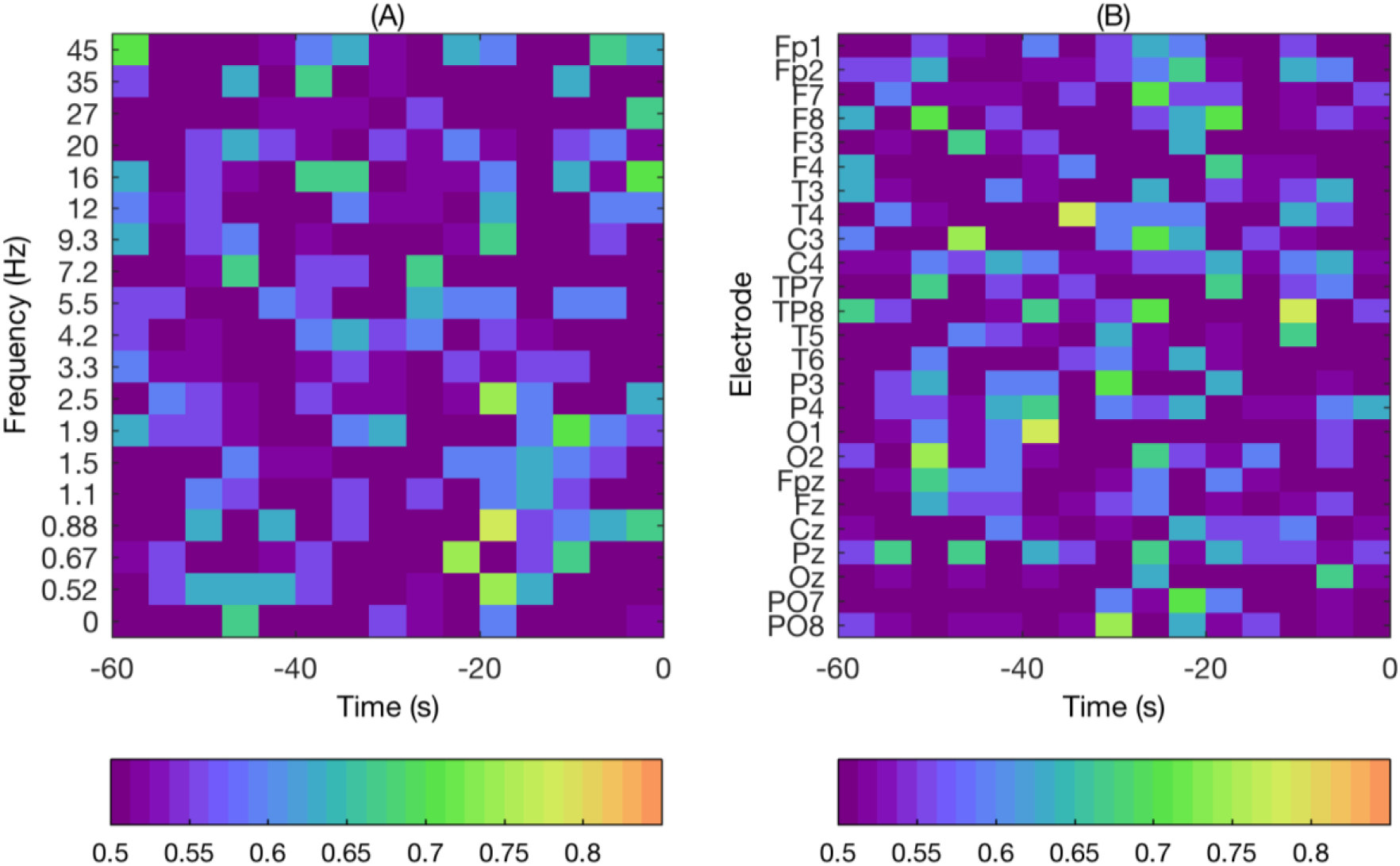
The time course of dreamfulness decodability for power spectral density features. Decodability is colour-coded according to the colour bars, with chance-level performance (0.5 accuracy) in dark purple. Decodability is shown for electrode-frequency power features: within each frequency (*A*) and within each electrode (*B*). The critical thresholds for statistical significance at uncorrected *p* ≤ .01, .001 and .0001 respectively are decodabilities .70, .78 and .85 (one-tailed Binomial test, *N* = 27).

### Summary

In all three analyses, we found no significant difference between dreamful and dreamless cases for the features we extracted, which included electrode-frequency spectral power, EMG and EOG activity. In our Bayesian modelling comparison, we found strong evidence against Siclari et al.’s (2017) high-frequency posterior hot zone effect. We also failed to replicate both Siclari et al. (2017) and Scarpelli et al.’s (2017) low-frequency effect to statistical significance.

## Discussion

The Dream Catcher test is a paradigm to test whether an understanding of the neural constituents of consciousness (i.e., experiences) is indeed genuine, by separating the measurement of brain activity from associated subjective reports. Furthermore, the test should be performed under the stipulation that the full contents of consciousness are generated by internal neural mechanisms operating spontaneously and independently of external stimulation. If a scientist can reliably predict the full contents of consciousness based only on observations of the neural activity and with no access to information on the external stimuli or subjective reports, then the test is passed.

We executed a simplified version of the Dream Catcher test with data from 9 participants, collected by the Data Team with an early night serial awakening paradigm. The Analysis Team’s task was to sort the EEG segments (i.e., 60-second polysomnograms preceding awakenings and interviews) into two groups: segments associated with dream reports vs segments associated with reports of non-dreaming. The Data Team evaluated the Analysis Team’s performance over five decreasing levels of blindness (Fig. 1): 1) all cases unlabelled, 2) cases paired by complementary dreamfulness conditions from the same participant, 3) cases labelled by common participant, 4) cases labelled by common dreamfulness condition and participant, and 5) cases labelled by common dreamfulness condition. The Analysis Team approached the classification task by clustering together quantitatively similar cases into two groups and then manually classifying those groups based on findings from the recent literature. The similarity metrics used in clustering were based on features extracted from power and location of EEG spectra, and EMG and EOG activity.

At all levels of blindness, the Analysis Team was unable to correctly classify between dreamfulness vs. dreamlessness with statistically significant accuracy; the best performance was achieved at Steps 2 and 3 with an accuracy of 59% (*p* = .44). Thus, the Analysis Team did not pass even this rudimentary form of the Dream Catcher test. Subsequently, several post hoc analyses were carried out to identify the source of failure. We went on to fail three more times at finding a difference between dreamful and dreamless cases using the fully-labelled data set with different paradigms: Bayesian model selection, MANOVA, and decodability. In particular, the model selection analysis failed to confirm key findings from the recent literature, while the MANOVA and decoding analyses found no effect amongst the selected range of features, which included EEG power spectra, and EOG and EMG activity.

### Explaining the Results

The result of this Dream Catcher experiment suggests that the neural correlates of dreaming consciousness reportedly found in the power spectra of the brain are not sufficient for bridging the explanatory gap of consciousness. Due to the challenging nature of our experimental setup, our failure to pass the Dream Catcher test was not in itself a surprising outcome. What was surprising, however, was our subsequent failure to replicate or confirm the findings of the studies from which we derived our original predictions. We address several possible explanations.

First, there were several methodological differences between the original studies and our attempted replication studies (see Table 4). On one hand, we had fewer participants than Chellappa et al. (*N* = 17), Esposito et al. (*N* = 11), Siclari, Baird et al. (*N* = 32), Scarpelli (*N* = 14), and Siclari, Bernardi et al. (*N* = 12), which exposed us to a greater risk of false negative results. On the other hand, we note that Scarpelli et al., Chellappa et al., and Esposito et al. ground their findings based on just circa 2–3 awakenings per participant, which would have led to more variable data within participants, in contrast to the 6 awakenings performed in our study. Data of our sample size theoretically allowed us to detect an effect of size *d* ≥ 0.78 for at least .8 statistical power and *α* ≤ .05 false positive probability, assuming two groups of *N* = 27 independent Gaussian-distributed samples each and no between-participant variability. More conservatively, if samples were averaged for each participant, we could detect effect sizes *d* ≥ 1.1, assuming *N* = 9 paired, Gaussian-distributed samples.

**Table 4.**
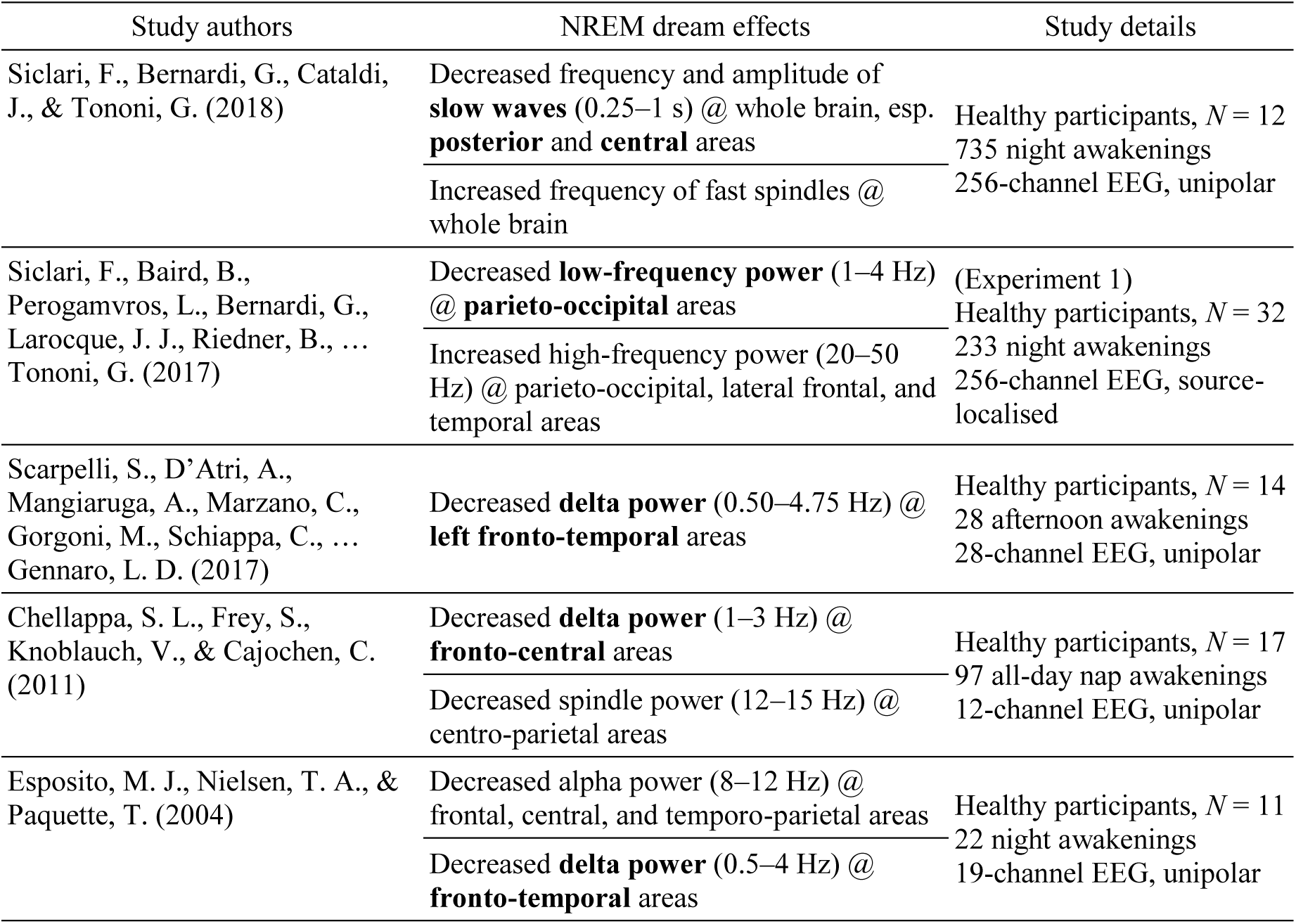
Comparison of similar studies

| Study authors | NREM dream effects | Study details |
| --- | --- | --- |
| Siclari, F., Bernardi, G., Cataldi, J., & Tononi, G. (2018) | Decreased frequency and amplitude of **slow waves** (0.25–1 s) @ whole brain, esp. **posterior** and **central** areas<br>Increased frequency of fast spindles @ whole brain | Healthy participants, $N = 12$<br>735 night awakenings<br>256-channel EEG, unipolar |
| Siclari, F., Baird, B., Perogamvros, L., Bernardi, G., Larocque, J. J., Riedner, B., ... Tononi, G. (2017) | Decreased **low-frequency power** (1–4 Hz) @ **parieto-occipital** areas<br>Increased high-frequency power (20–50 Hz) @ parieto-occipital, lateral frontal, and temporal areas | (Experiment 1)<br>Healthy participants, $N = 32$<br>233 night awakenings<br>256-channel EEG, source-localised |
| Scarpelli, S., D'Atri, A., Mangiaruga, A., Marzano, C., Gorgoni, M., Schiappa, C., ... Gennaro, L. D. (2017) | Decreased **delta power** (0.50–4.75 Hz) @ **left fronto-temporal** areas | Healthy participants, $N = 14$<br>28 afternoon awakenings<br>28-channel EEG, unipolar |
| Chellappa, S. L., Frey, S., Knoblauch, V., & Cajochen, C. (2011) | Decreased **delta power** (1–3 Hz) @ **fronto-central** areas<br>Decreased spindle power (12–15 Hz) @ centro-parietal areas | Healthy participants, $N = 17$<br>97 all-day nap awakenings<br>12-channel EEG, unipolar |
| Esposito, M. J., Nielsen, T. A., & Paquette, T. (2004) | Decreased alpha power (8–12 Hz) @ frontal, central, and temporo-parietal areas<br>Decreased **delta power** (0.5–4 Hz) @ **fronto-temporal** areas | Healthy participants, $N = 11$<br>22 night awakenings<br>19-channel EEG, unipolar |

On the technical level, Siclari, Baird et al.’s sleep study differed from ours. Whereas they used a 256-channel high-density EEG system, we were limited to a 29-channel system; and instead of dipole current source modelling, we used Perrin’s method of estimating scalp current source density by taking the Laplacian of fitted spherical splines (Perrin, Pernier, Bertrand, & Echallier, 1989). Our replication of Siclari et al.’s hot zone measurements was therefore less precise. Nonetheless, such precision was not apparently essential to detecting an effect, as in Experiment 3 of their study, Siclari et al. were also able to successfully detect and predict the occurrence of dreaming vs. non-dreaming sleep utilising plain surface electrodes. Our data were more similar to Scarpelli et al.’s sleep study: they used 28 scalp electrodes with unipolar referencing for their analysis.

Another explanation for our results might have to do with extraneous variability in our data. Clustering works by grouping data according to their relative positions in feature space, but if these positions are influenced more by irrelevant factors or random noise than the relevant effect, then clustering would produce noisy or incorrect results. This was readily apparent from the result of Step 1, where clustering in fact grouped cases together by participant identity and not by dreamfulness (Figs. 3*B* & 9). However, even though participant identities were balanced from Step 2 onwards using pairing information, the classification accuracy did not significantly improve. This remained true even after removing independent components of EEG time series that clustered contrarily to pairing and participant information in Steps 3 and 4. Furthermore, the lack of an interaction effect found in the MANOVA meant that there was no evidence that the effect of dreamfulness might have varied between participants either. If our analysis failed to find an effect due to the presence of irrelevant factors or noise, it is unlikely due to only inter-participant variability. Assuming that an effect does exist, we would require a larger sample size to measure it.

Lastly, we defined dreams as any experiences occurring during sleep—an extremely simple and broad definition in comparison to more fine-grained conceptual frameworks of sleep experience (see Windt et al., 2016). In fact, all NREM sleep dreams included in our study were static (i.e., lacking change or temporal progression), whilst other studies might treat only complex, dynamic and temporally progressing states as genuine instances of dreaming (Hobson et al., 2000; Hall, 1953; Nielsen, 2000; Revonsuo, 2006). It is possible that we failed to classify dreamful and dreamless NREM episodes, or find similar patterns of results as in previous studies, due to differences in how each study defined dreaming and its subtypes. Chellappa et al. (2011) asked participants “How much did you dream?” and identified a given report as dreamful when the answer was “greatly”, “fairly”, or “little”, likely including both static and dynamic dream reports in the study sample. Siclari et al. (2017) categorised both perceptual and non-perceptual mentation reports as cases of dreaming as long as participants “had been experiencing anything”. Scarpelli et al. (2017) selected dreams with narrative and temporal properties (Foulkes & Schmidt, 1983), and disregarded dreams without recall. On the other hand, dreamless sleep was identified similarly by all these studies as reports of having no experience.

### The Relation Between Dreamfulness and the Depth of NREM Sleep

Even though a number of studies reported that NREM sleep dreaming is associated with a decrease of delta power over differing locations in the brain (Chellappa et al., 2011; Esposito et al., 2004; Siclari et al., 2017, 2018, 2014), we were not able to replicate this finding. While this could be due to an unidentified confound in our experimental design, data, or analysis, it is also possible that delta power decrease is not a genuine correlate of dreaming. But here, we propose another possible explanation: that delta power may simply reflect the depth of NREM sleep rather than dreaming consciousness.

NREM Stages 2, 3 and 4 have been classically delineated by the proportion of delta waves within the course of a 20- or 30-s epoch (Rechtschaffen & Kales, 1968). Stage 2 sleep was primarily defined by the presence of sleep spindles and up to 20% prevalence of delta waves; Stage 3 sleep was defined for having 20–50% delta waves; and epochs with >50% delta waves were scored as Stage 4 sleep. By these definitions, there is a huge variance in the amount of delta waves not only between sleep stages but also within a given NREM stage. This has become even more amplified by the sleep scoring guidelines introduced by the American Academy of Sleep Medicine (Iber, Ancoli-Israel, Chesson, & Quan, 2007) with the merging of NREM Stages 3 and 4 into the N3 stage of “slow wave sleep” (delta waves dominating 20-100% of the time). Importantly, dream recall is already known to decrease from Stage 2 to Stage 3 and subsequently from Stage 3 to Stage 4 (Foulkes, 1982; Moffitt, 1982; Noreika et al., 2009; Pivik, 1971; Pivik & Foulkes, 1968). As these stages are delineated by the amount of delta waves, it is plausible that the correlation between dream recall and delta power might be confused for the correlation between dream recall and NREM depth. Thus, the previously reported association between dreaming and EEG delta power may reflect a simple correlation between dream recall and the depth of NREM sleep (within each Stage) rather than intrinsic neural mechanisms of dreaming.

As an example, Siclari et al. (2017) were able to predict dream recall in real time by awakening participants once delta power decreased below an individual spectral threshold (in addition to a gamma increase). Arguably, such awakenings yielded dream reports that took place during relatively shallow NREM sleep, whereas in deeper NREM sleep, we would expect to have more dreamless sleep reports. Such an uneven association between the presence of dream experience and the depth of sleep even within a given NREM sleep stage is possible when all reports of dreaming and dreamless sleep are used for EEG analysis. In our study, the confounding factor of the sleep stage (or depth) was minimised by using only a small matched subsample of data (*N* = 27 + 27) from a larger pool of awakenings (*N* = 294), and making sure we had equal numbers of N2 and N3 sleep in dreamful and dreamless conditions from the same participant. This reduced a distribution bias of more dreamless reports at deeper stages of sleep.

### Pros and Cons of the Dream Catcher Paradigm

We have found, through the course of our experiment, that the constraints imposed by the Dream Catcher paradigm force the researchers to focus their efforts on a single determination of the data. This mindset is considerably different to the status quo of scientific research nowadays, where one can pursue multiple avenues of investigation, sequentially or simultaneously, and then deal no further with those yielding nonsupporting results. Those failed avenues of investigation end up being incompletely explained and typically remain unpublished. By contrast, the Dream Catcher test permits investigating only a single avenue, and the researcher must address its results whether they support the hypothesis or not. It is similar in spirit to the Registered Reports format for science publishing (see Chambers, 2013 for an exposition). In addition, blinding the data removes bias; unlike in common practice where the results are always known, researchers that pass the Dream Catcher test do so blindly and based on genuine understanding. The Dream Catcher paradigm encourages not only good science, but also a critical assessment of the reliability of past findings.

This paradigm is not, however, without costs. Compared to the testing of multiple hypotheses, the data for a Dream Catcher test—in principle—can only be used once per hypothesis, which is relatively inefficient. Even if there were multiple Analysis Teams working in parallel, there would be no way to ensure that they each tested different hypotheses—because any such attempt would undermine the independence of the teams and could reintroduce bias. We addressed this inefficiency problem somewhat in our design by re-evaluating performance iteratively with a gradual removal of blindness. We hoped this would give us insights into the level of information required for successful classification.

Following the completion of this experiment, we once again address the inefficiency problem and propose that multiple hypotheses could be validly tested within this paradigm through the use of unsupervised machine learning algorithms; for example, by setting up separate algorithms to test each hypothesis and disregard information that might otherwise produce experimenter bias. In our study, we started with the assessment of several families of features for clustering consistency, but ultimately submitted our answers based on just one of them. If we had allowed for the submission of multiple answers in parallel for each set of features, we would have been able to broaden the scope of our study. The caveat, however, is that once again the researcher may be tempted to neglect nonsupporting results at the next step.

With regard to the use of predefined algorithms to achieve unbiased results, clustering, as used by the Analysis Team, is not the only method. The rationale for using clustering was motivated by the desire to discriminate cases based on several measures simultaneously in order to maximise performance; however, it may well be the case that one aims to test a single measure at a time. In that case, one could resort to very simple procedures. For example, for two equal-sized classes, one could use a median split; for two classes without assuming equal sizes, an automatic thresholding algorithm; and for multiclass classification, *k*-means clustering coupled with a classification heuristic. Researchers should determine for themselves the best method to represent the hypothesis they wish to test.

### Future Avenues of Investigation

The data analysed here constitute only a portion of the total number of awakenings (294) performed and recorded in our sleep study. Now that we have completed the Dream Catcher experiment with all its self-imposed restrictions, we can further investigate the replicability issue with the data we did not include. This may well reveal that those unreplicable effects do in fact exist, but they did not survive our precursory data selection process: the balancing of NREM stages and degree of dreamfulness.

Additionally, we can check other effects reported in past studies within our expanded data set. Contemporary theories of consciousness suggest that connectivity or integration in the brain is necessary for the emergence of consciousness. The studies we have addressed here, as well as ours, did not consider such features; all features were measures of univariate data only operating on one channel at a time. To measure connectivity, we must employ features that operate on multivariate data. Phase coherence and cross-correlation, as we have mentioned in the introduction, are well-known examples. More sophisticated measures have been proposed for quantifying consciousness (Barrett & Seth, 2011; Oizumi, Amari, Yanagawa, Fujii, & Tsuchiya, 2016; Tegmark 2016; Schartner et al., 2015; Kim et al., 2018), taking inspiration from the integrated information theory of consciousness (Oizumi, Albantakis, & Tononi, 2014; Tononi, 2004, 2008). These measures are all necessarily multivariate in nature. It is possible that the true correlates of dreaming consciousness in NREM sleep are to be found in multivariate features rather than univariate ones. We intend to investigate this possibility in future work.

## Acknowledgements

William Wong was supported by an Australian Government Research Training Program Scholarship. Valdas Noreika was funded by the Signe and Ane Gyllenberg Foundation. Jennifer Windt was funded by an Australian Research Council Discovery Early Career Researcher Award (DE170101254). Katja Valli was funded by the Academy of Finland (project 266434). Naotsugu Tsuchiya was funded by an Australian Research Council Future Fellowship (FT120100619) and Discovery Project grants (DP130100194, DP180104128, DP180100396). We thank Mila Oravecz and Lisa Svartsjö for the contribution to the data acquisition and dream content analysis, and Matti Grönroos and Petri Vähä-Pesola for the development of statistical models to assess performance of the Analysis Team. PSG data used in the Dream Catcher experiment are available upon request.

## Supplementary Document 1

### Sleep Data Collection Methods

#### Early night serial awakening

The experimental data were collected during four non-consecutive experimental nights, which took place between 10 p.m. and 6 a.m., with at least three nights between any two sessions. The participants were requested to avoid any stimulants, such as caffeine (6 hours prior to the experiment) or over-the-counter medication and alcohol (24 hours prior to the experiment), which was controlled for by asking participants to fill out a questionnaire before each session. The participants slept in a separate room and were observed through a video camera. The team constantly monitored their PSG on a computer screen and communicated with participants through a digital sound system device.

During each experimental night, participants were awakened during the first 3–4 hours on average 8.17 times (*SD* = 1.24, range 6–11), yielding a total of 294 awakenings (9 participants × 4 nights). This early night serial awakening protocol was shown to be an efficient paradigm for collecting large samples of dream reports while maintaining the stability of EEG spectral power measurements throughout the session (Noreika et al., 2009).

The experimenters awakened the participants by playing a beep sound after confirming through online monitoring of PSG that they had been in Stages 2 or 3 of NREM sleep for at least 1 to 3 minutes. Post-hoc examination of the EEG confirmed that most of the awakenings took place during NREM Stages 2–3 (87%) over the 1-minute period before awakening. Given that serial awakenings from NREM sleep acted as selective REM sleep deprivation, episodes of sleep-onset REM occasionally intruded into the normal progression of sleep stages (13%), in which case participants were awakened as usual. Data from REM awakenings were omitted from this study.

Before the experimental session, the participants were instructed in advance that, immediately after being awakened by the sound signal, they were to give a free oral report of “everything that was going through their mind before awakening”; the procedure was practised during the adaptation night. This instruction was not repeated again after the individual awakenings throughout the night, as a non-prompted free dream report was expected to interfere as little as possible with the very delicate process of remembering their experiences. If the experimenter judged that a free report from the participant contained any pre-awakening thoughts or perceptual experiences, the participant was further examined with a pre-recorded set of 21 questions played on a computer via the sound system (see Supplementary Document 2 for detailed instructions given to participants). The questions included inquiries about objects, feelings, self, and the subjective duration of the dream. Several questions aimed to reinforce recall. If the experimenter judged that a free report from the participant contained no pre-awakening thoughts or perceptual experiences, it was followed by three questions regarding the subjective certainty of dreamless sleep. In cases where these three questions prompted the participant to remember any experiences, they were presented with the set of 21 questions concerning the contents of their dream. If necessary, an unstructured interview was conducted at the end to clarify unclear or ambiguous parts of the report. The experimental night was ended at the wish of the participant or when the number of awakenings was satisfactory. All reports, questionnaire answers, and interviews were recorded on a computer.

#### EEG acquisition

The EEG montage included 21 electrodes placed according to the standard 10-20 system and 4 additional electrodes (TP7, TP8, PO7, PO8) placed according to the 10-10 system (Oostenveld & Praamstra, 2001). All these electrodes were referenced to the right ear mastoid, and the ground electrode was placed on the temple. In addition, two bipolar EOG electrodes were placed near the lateral canthus and the lower eyelid to measure eye movement, and a pair of bipolar electrodes was placed on the mentalis and submentalis muscles to record chin EMG. Recordings were carried out on a SynAmps amplifier and NeuroScan (4.1.1) data acquisition software at a 2,000 Hz sampling rate. EEG data were saved in the frequency band of 0.05–100 Hz (with a 50 Hz notch filter and a gain of 1,000), EOG data in the band 0.05–30 Hz (gain 1,000), and EMG data in the band 5–500 Hz (gain 2,500). Electrodes were silver chloride and attached to the skin using Grass EC2 electrode cream.

#### Content analysis of post-awakening reports

The post-awakening reports and interviews were transcribed for content analysis, consisting of the following two stages.

First, all reports were divided by two independent raters (Master students in psychology) into four categories: 1) dreamless sleep, 2) white dream, 3) uncertain, and 4) dream (following Dement, 1955). Reports were scored as dreamless if the participant was confident they had no experiences right before awakening. Reports were scored as white dreams if the participant strongly felt they had had some experiences right before awakening, but could not recall any specific content. Reports were scored as uncertain if the participant was unsure whether they had been dreaming or had dreamless sleep right before awakening. Finally, reports were scored as dreams if participants reported any experiences (e.g., perceptions in any sensory modality, sensations, thoughts, feelings, and emotions). Inter-rater reliability for the 4-way categorisation of reports was 94% (*κ* = .92).

Second, the dream reports were further categorised by the same two independent raters using Orlinsky’s Modified Scale for Perceptual Complexity of Dreams (Noreika et al., 2009; Orlinsky, 1962). This scale consists of 7 perceptual complexity categories, ranging from “1=Participant remembers a specific topic but in isolation: a fragmentary percept, unrelated to anything else” to “7=Participant remembers a long, detailed dream in which the whole scene is replaced by other scenery more than once”. Categories 1–4 depict static dreams that lack any change or temporal progression, whereas categories 5–7 depict dynamic dreams that contain a change of at least one perceptual experience (Noreika et al., 2009). The inter-rater reliability for the 7-way categorisation of dream reports was 83.8% (weighted *κ* = .89), with most errors consisting of reports assigned to adjacent complexity categories. During both stages of content analysis, the two raters discussed scoring disagreements until agreement was achieved, or—in a few cases—a third rater (author VN) was asked to judge which of the two suggested categories was more accurate.

#### Sleep scoring

Sleep stages were manually scored by authors VN and KV, and each 1-minute EEG recording was scored as three 20-second epochs. Initially, the scorers agreed on 76% of the epochs. For the remaining 24%, the scorers discussed them until they reached a consensus. Scoring criteria defined by Rechtschaffen and Kales (1968) were followed, which allowed for a more fine-grained consideration of the amount of delta waves during slow wave sleep than the most recent sleep scoring guidelines (Berry et al., 2012).

## Supplementary Document 2

### Dream Report Interview Procedure

#### Preface

The following is an English translation of the original dream report interview procedure instructions in Finnish, used by the Data Team to conduct the post-awakening dream report interviews. Some examples of the dream reports are given in Supplementary Document 3.

#### Introduction

The aim of this research project is to investigate the neural correlates of dream experiences during NREM sleep. You will spend five nights in the sleep laboratory: one adaptation night during which we will wake you up four times and you can practise dream reporting, and then four experimental nights during which we will wake you up several times (5–10) from early night sleep, both from light sleep and from deep sleep, and request you to report any dream experiences you might have had. Due to the awakenings, your total sleep time will be shorter than usual, and you may feel tired in the morning and the next day. If you wish to terminate the experiment, you can do so at any time.

#### Wake-up procedure and reporting instructions

When you have fallen asleep and are in a specific sleep stage, we will wake you up with a sound signal. When you hear the sound signal and wake up, try to stay still and calm, as this way your dream recall is least compromised. We will not ask any questions at this point. Your task is to give us a free recall report: tell us whether you remember having a dream experience during sleep and what the experience was like. Tell us every detail you can remember. By dream experience, we mean every image or thought you had in mind just before waking up.

We do not expect you to recall a dream every time we wake you up. For this research project, it is of utmost importance that you honestly report what you remember, and also honestly report if you don’t remember anything.

When you have recited everything you recall of the dream, we will ask further questions. The questions have been pre-recorded, and we will ask all participants the same questions. Therefore, some of the questions may not feel relevant in the context of your dream. Regardless, try to answer as well as you can, and be honest. If you cannot answer a question, let us know that you cannot remember or tell more about the requested detail. You also have the right to protect your privacy: If you do not wish to share a specific detail of your dream, tell us, and we will not ask further questions about this detail.

If you do not recall any experiences, we will also ask some further questions. Again, the questions have been pre-recorded, and we will present the same questions to all participants.

If we wish to know more about your dream than the recorded questions reveal, we may also ask you some specific, unique questions after the recorded questions.

We will record all your answers to be able to later assess them.

#### The dream interview and how to answer the questions

##### If you do not recall a dream

If, after the sound signal, you report that you do not remember any dream experiences, we will ask you the following questions (N1–N3):

N1) How certain, on a scale from 1 to 5, you are that you did not have any dream experiences?

1 = very uncertain

2 = quite uncertain

3 = cannot say

4 = quite certain

5 = very certain

N2) Do you have the impression that you had a dream experience but cannot recall any specific content?

If you have the impression that you were dreaming, but cannot recall any content at all, answer yes. If you are certain that you did not have any dream experiences, answer no.

N3) What is the last thing you recall?

Tell us the last thing you remember. You don’t need to know whether this was from before falling asleep or whether it occurred while asleep.

##### If you recall a dream

After you have heard the sound signal and you have reported your dream in as much detail as you can remember, we will ask the following questions (P1–P17):

P1) Did you have the experience just before waking up?

Your task is to estimate when the dream took place. Did the sound signal wake you up in the middle of the experience? Answer yes if you feel that this was the case. If you feel that the dream took place earlier and you had no experiences at the time you woke up, answer no.

P2) How certain, on a scale from 1 to 5, are you that the dream experience took place just before you woke up?

1 = very uncertain

2 = quite uncertain

3 = cannot say

4 = quite certain

5 = very certain

P3) Describe the setting where the dream took place.

Your task is to describe the environment in which the dream events occurred. What was the setting like?

P4) List all the objects in your dream.

Your task is to list all the objects you recall appearing in the dream. No object is too irrelevant to be named.

P5) List all the sounds you heard in your dream.

Your task is to list all the sounds you recall hearing in the dream, such as speech, traffic, or music.

P6) List all the characters in your dream, including also those other than human.

Your task is to list all the human and other animate characters you recall appearing in the dream (animals, fantasy figures, etc.).

P7) Describe all the emotions and feelings and moods you experienced in the dream.

Your task is to describe all the emotions, moods, and feelings you recall having in the dream (e.g., happiness, joy, love, affection, sadness, hate, anger, fear, anxiety, disgust, surprise).

If you did not have any emotional experiences in the dream, report that no emotions were present. Then we will move on with the questions. If you recall having experienced at least one emotion in the dream, we will ask you two additional questions (P7a–P7b):

P7a. What dream event were the emotions related to?

Describe the event, thought, action or character the emotion was related to or in response to. If you had several emotional experiences, list each, and describe the element each emotion was related to.

P7b. Estimate the intensity of the emotion on a scale from 1 to 5.

Your task is to estimate the intensity of the emotion. If you experienced several separate emotions, please state the name of the emotion first and then estimate its intensity.

1 = very low

2 = quite low

3 = moderate

4 = quite intense

5 = very intense

P8) List all the changes that took place in your dream.

If an object changed into another, if something changed in the setting in which the dream took place, or if a new thought or emotional state appeared, list those.

P9) Were you present or embodied in the dream?

Your task is to tell us whether you were present in the dream and participating in the dream events. If you had a body in the dream, answer yes. If you experienced the dream as a bodiless spectator or an outside observer, like watching a movie, answer no. If you answer yes, we will ask you two additional questions (P9a–P9b):

P9a. Describe your dream body.

Describe what kind of a body you had in the dream. Was your dream body identical or similar to your physical body? If it was different to your physical body, how was it different? Aim to describe your dream body in as much detail as possible.

P9b. Were you an active participant in the dream or just an observer?

Describe whether you participated actively in the dream events from an embodied perspective (e.g., talking, thinking, moving, looking around) or whether you were merely a passive spectator or observer.

P10) How clear was the perceptual quality in your dream, on a scale from 1 to 5, compared to how you perceive the world when you are awake?

Your task is to estimate how well the perceptual quality of your dream experience matched the perceptual quality of normal waking experiences. Evaluate how clear the visual, auditory, olfactory, gustatory, tactile and kinaesthetic perceptions and sensations were compared to comparable experiences during wakefulness:

The quality of the perceptions and sensations was

1 = very vague and obscure

2 = quite vague and obscure

3 = almost as clear and defined as during wakefulness

4 = equally clear and defined as during wakefulness

5 = extremely clear and defined, more so than during wakefulness

P11) Estimate the duration of your dream experience.

Estimate how many minutes or seconds your dream experience lasted. If you cannot make an estimation, guess.

1 = about few seconds

2 = about half a minute

3 = about one minute

4 = longer than three minutes

5 = longer than ten minutes

6 = longer than half an hour

P12) How certain are you of your answer to the previous question (P11)?

1 = very uncertain

2 = quite uncertain

3 = cannot say

4 = quite certain

5 = very certain

P13) How quick was the passage of time in your dream?

Your task is to evaluate the subjective experience of passage of time in your dream. How quickly did time seem to be passing compared to how time passes during wakefulness? How fast did the objects or characters move or change in your dream compared to wakefulness?

Time passed:

1 = much slower

2 = a bit slower

3 = comparably to waking experiences

4 = a bit faster

5 = a lot faster

P14) How certain are you of your answer to the previous question (P13)?

1 = very uncertain

2 = quite uncertain

3 = cannot say

4 = quite certain

5 = very certain

P15) What were the factors that allowed you to estimate the passage of time in your dream?

P16) Estimate how long you slept.

Estimate how many minutes or hours you had slept (since the last awakening) before we woke you up.

P17) Do you remember anything else?

If you recall anything else that you have not reported yet, please report what these experiences were.

#### Postface

The following are summary statistics to some answered questions that the reader might find relevant, counted over the 27 dreamful reports used in the Dream Catcher experiment.

P2) How certain, on a scale from 1 to 5, are you that the dream experience took place just before you woke up?

85% very certain or certain

7% cannot say

7% quite uncertain

P10) How clear was the perceptual quality in your dream, on a scale from 1 to 5, compared to how you perceive the world when you are awake?

15% very vague and obscure

30% quite vague and obscure

33% almost as clear and defined as during wakefulness

19% equally clear and defined as during wakefulness

4% extremely clear and defined, more so than during wakefulness

P11) Estimate the duration of your dream experience.

22% about few seconds

33% about half a minute

26% about one minute

11% longer than three minutes

7% longer than ten minutes

0% longer than half an hour

## Supplementary Document 3

### Examples of Dream Reports

The following are examples of subjects’ dream reports after being awakened, translated from the original Finnish reports. All reports were scored as static in Orlinsky’s scale (score 1–4). Most were composed of several interconnected perceptions (score 3–4), and most also included a unified background scene (score 4).

### Subject 4, 3rd experimental night, awakening 8

“There was a dry open field, and a dog on an agility track. On the track, there was a tube and hurdles. I was uncertain what I was supposed to do there.”

Orlinsky score = 3

### Subject 5, 1st experimental night, awakening 2

“I dreamt about a green car parked on a street. I saw the car from the side; it was bright green, and I heard the engine running.”

Orlinsky score = 3

### Subject 8, 4th experimental night, awakening 3

“I saw a cafeteria patio on a street.” Orlinsky score = 1

### Subject 11, 3rd experimental night, awakening 3

“I was in a harbour, and I think I had loaded something in some boat. It looked like a Greek harbour, with several boats tied to poles. The sun was shining; on the other side was the sea; and on the other, a forest or something. I think my boyfriend and friends were there with me.”

Orlinsky score = 4

### Subject 15, experimental night 1, awakening 3

“I dreamt I was in this lab, in this room, and about the electrodes in my head. This bed was there, and someone was putting on the electrodes. In the background, there was a sound of someone talking.”

Orlinsky score = 4

### Subject 16, experimental night 2, awakening 5

“I was at a party—I think it was my graduation party. We were outdoors in a yard; there was grass, and tables with white table cloths; some dishes and a cake on the table. There were a few people there; I heard them talking.”

Orlinsky score = 4

## Supplementary Document 4

### Analysis Team Briefing Document

#### Preface

The following is the briefing document provided by the Data Team to the Analysis Team prior to the commencing the Dream Catcher experiment.

#### Dream Catcher Experiment: Gradual removal of blindness

CRG, Turku

17-03-2008

#### Data and general procedure

Our data set comes from 9 subjects with 3 dream and 3 dreamless sleep reports from each: in total, there are 54 EEG recordings.

The data will be provided to you in 5 steps from the most complex to the easiest one; a following step will always take place only when a previous one has been completed and its success rate has been evaluated. With this kind of procedure, we will get several different decisions where the chance level of successful rating can be evaluated. Consequently, we will be able to see whether the EEG contain information about dreaming, and at what level that information is embedded (e.g. at the level of individual pairs of data cases, at the level of single subjects, or at the level of conditions).

Actual EEG data will be provided only for the Step 1 analysis. During Steps 2-5, only new file names will be given, which, in addition to new information, will always retain the previous ID codes of cases, pairs, subjects, etc.

#### Glossary

Case: a single 1 min pre-awakening EEG recording of either dream or dreamless sleep (provided in Step 1).

Pair: two 1 min pre-awakening EEG recordings of the same subject; one of them is dream and another one dreamless sleep recording (provided in Step 2).

Subject: information about each subject providing 3 pairs of EEG cases (provided in Step 3).

Group: within each subject, two groups of 3 EEG cases are formed; one group consists of 3 dream cases and another group consists of 3 dreamless sleep cases (provided in Step 4).

Condition: the whole data set is divided into two condition with 27 cases in each; one condition has only dreamless sleep cases and another – only dream cases (provided in Step 5).

#### Step 1: 54 *cases* are provided

##### DATA

Data is provided as 54 coded EEG cases: 27 dream and 27 dreamless sleep recordings (see Fig. 1). You remain blind to which EEG case originates from dreaming and which from dreamless sleep. Each subject will provide 6 cases – 3 dreams and 3 dreamless, but information on which cases belong to the same subject won’t be given at this stage.

**Figure 1.**
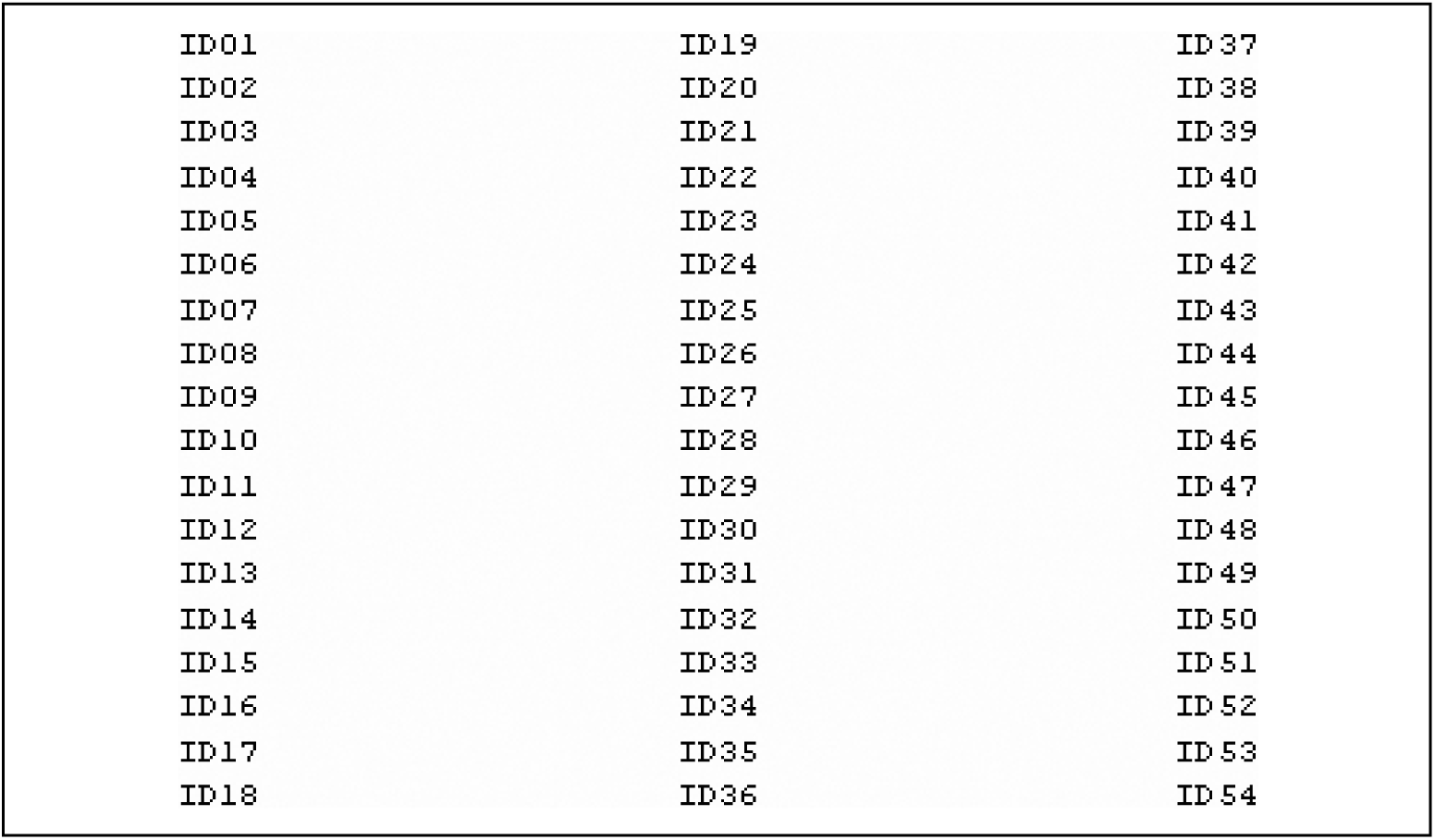
Example of 54 coded files names for the Step 1 analysis.

##### TASK

Your task is to decide which EEG case is “dreaming” and which is “dreamless sleep”, and to classify the cases better than at the chance level. Decisions must be based on the analysis of individual cases!

##### RESULTS

At least 38 cases out of 54 should be correctly identified in order to succeed above chance level. Also, each research group is expected to provide methodological rationale on what basis the decision was made, that is, which features in the EEG predict dreaming or non-dreaming.

Feedback will be given about overall success rate only, not about correct classification of individual cases.

#### Step 2. Single *pairs* of dream and dreamless sleep are provided

##### DATA

Data is provided in 27 coded EEG pairs, with 1 dream and 1 dreamless case in each pair (see Fig. 2). You remain blind to which EEG case originates from dreaming and which from dreamless sleep. Both cases of a single pair come from the same subject. Each subject will provide 3 pairs, but information on which 3 pairs belong to the same subject won’t be given.

**Figure 2.**
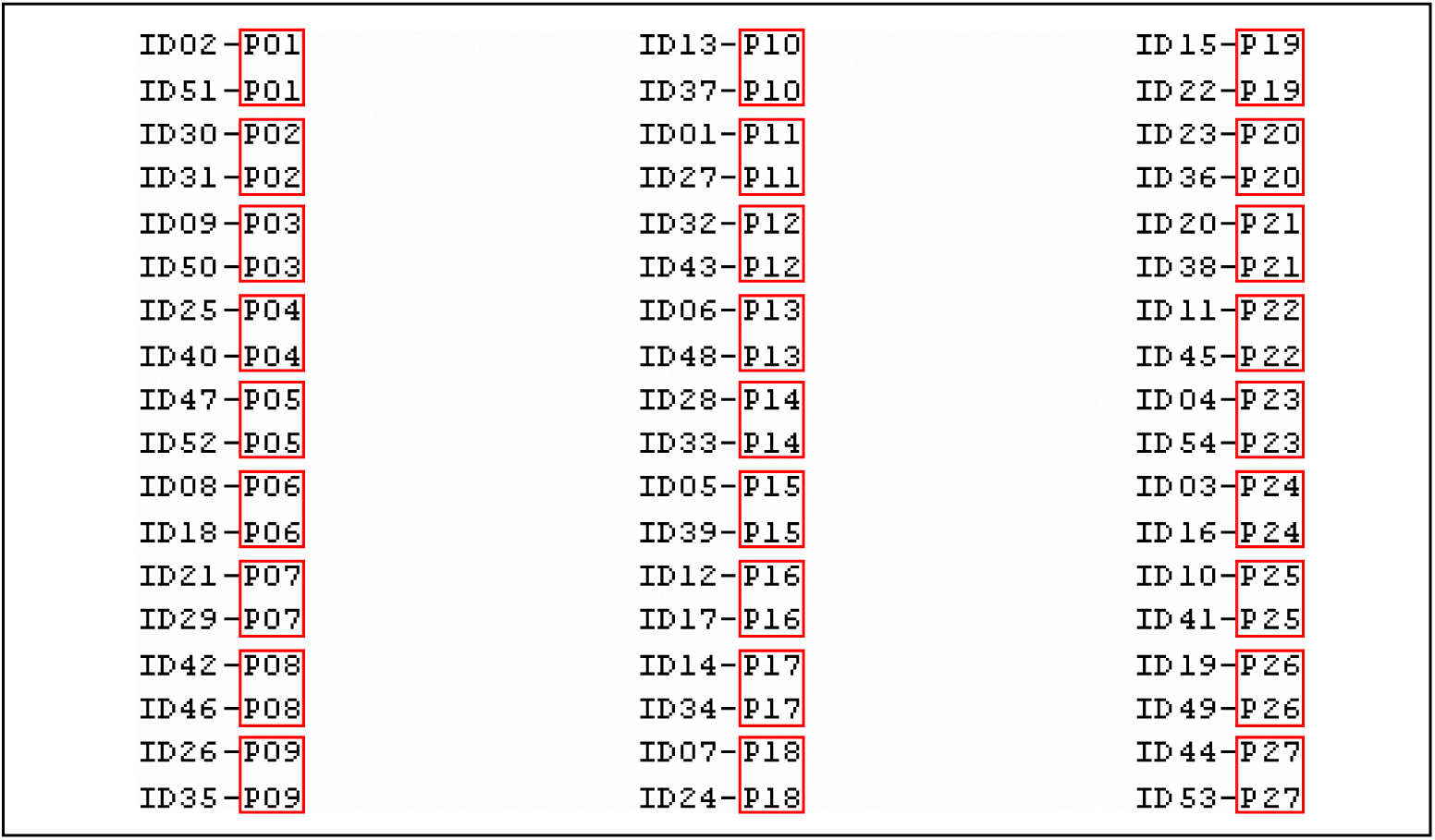
Example of 54 coded files names for the Step 2 analysis.

##### TASK

Your task is to decide which case in each EEG pair is “dreaming” and which is “dreamless sleep”, and to classify the cases better than at the chance level. Decisions must be based on the analysis of individual pairs, i.e. independent from other pairs!

##### RESULTS

At least 18 pairs out of 27 should be correctly identified in order to succeed above chance level. Also, each research group is expected to provide methodological rationale on what basis the decision was made, that is, which features in the EEG predict dreaming or non-dreaming.

Feedback will be given about overall success rate only, not about correct classification of individual pairs.

#### Step 3: *Subject* information for pairs is provided

##### DATA

Data is provided in 27 EEG pairs with 1 dream and 1 dreamless case in each, but it is not indicated which is a dreamless and which is a dream case (see Fig. 3). This time, however, 3 pairs of each subject are identified, so that comparison of pairs from the same subject is possible.

**Figure 3.**
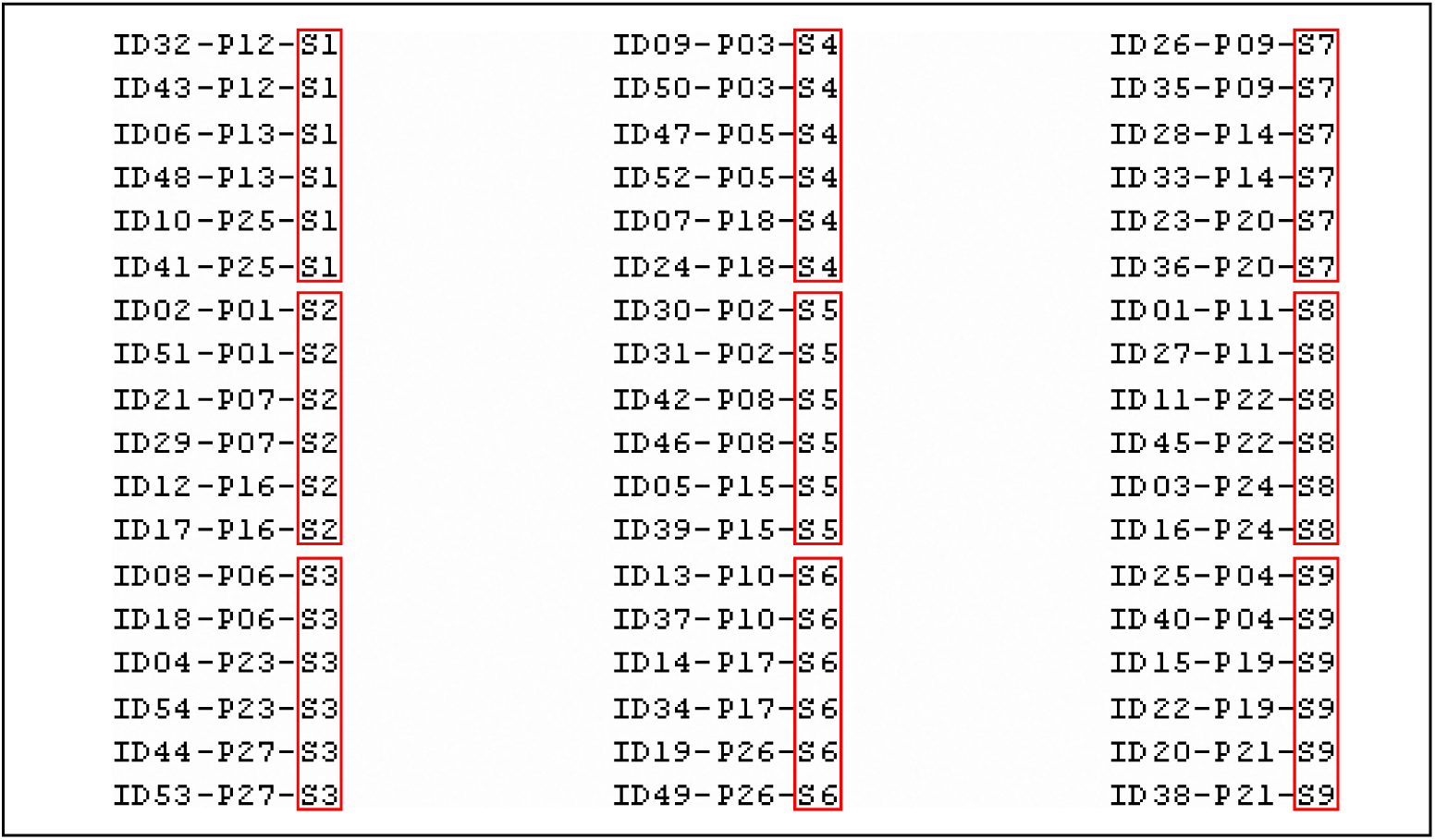
Example of 54 coded files names for the Step 3 analysis.

##### TASK

Your task is to decide which case in each pair is “dreaming” and which “dreamless sleep”, and to classify the pairs higher than at chance level. Still, decision should be done pairwise only, as no averaging between pairs is possible at this stage (averaging based on correct pooling of the type of cases is made possible at the next stage). At this stage, it is possible to compare EEGs of different individuals, and different pairs within the same individual, which should reduce variability in the data analysis.

##### RESULTS

At least 18 pairs out of 27 should be correctly identified in order to succeed above chance level. Also, each research group is expected to provide methodological rationale, on what EEG basis the decision was made, that is, which features in the EEG predict dreaming or non-dreaming.

Feedback will be given about overall success rate only, not about correct classification of individual pairs.

#### Step 4: *Groups* of cases within each subject are provided

##### DATA

Within each subject, two groups of different experimental conditions are provided (see Fig. 4). One group contains 3 EEGs prior to awakenings leading to dream report, and another group has 3 EEG cases from dreamless sleep of the same subject. However, it won’t be revealed, which is which. You will get 18 such groups coming from 9 subjects, yet, no identification across all subjects and their groups will be provided.

**Figure 4.**
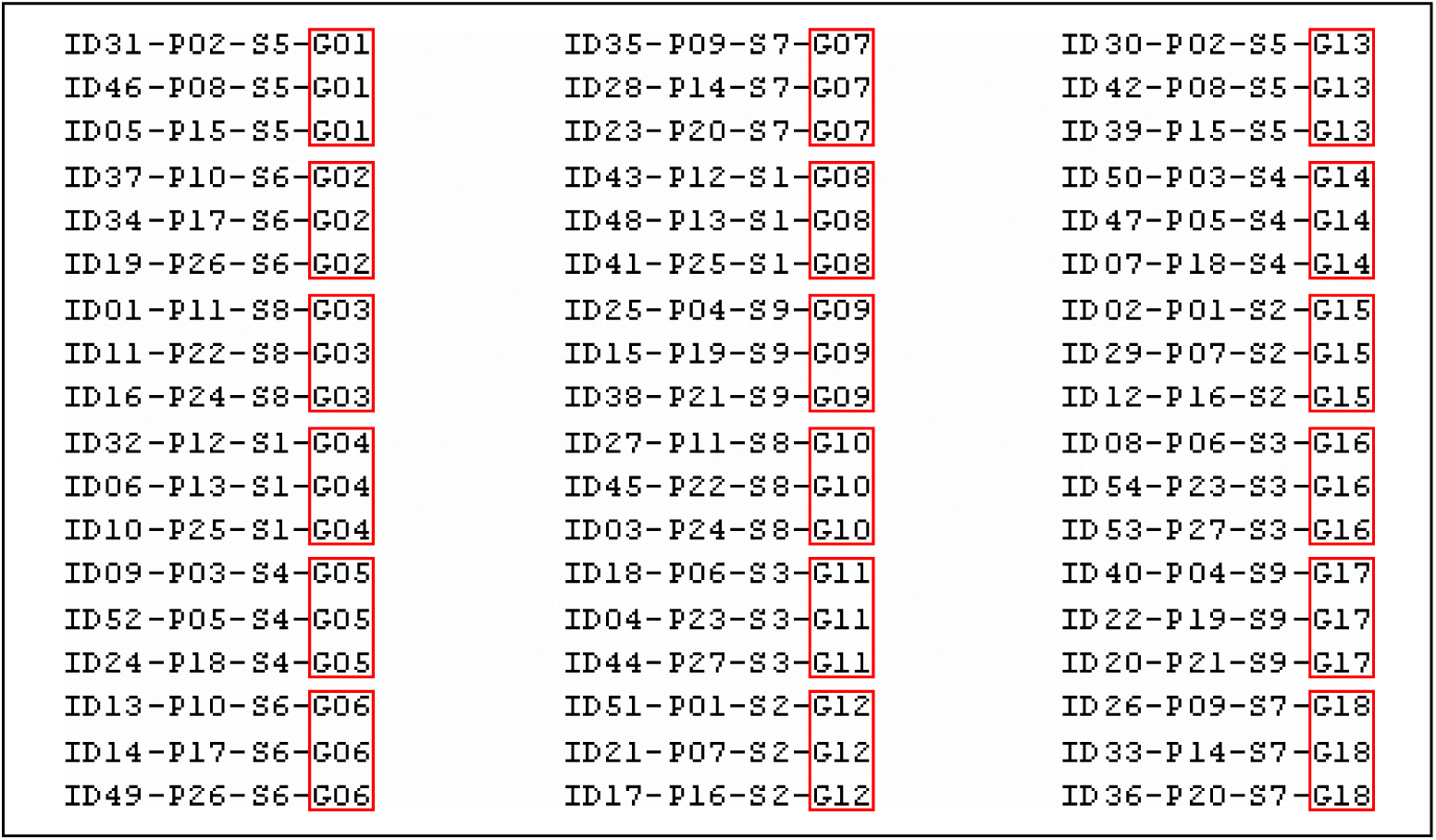
Example of 54 coded files names for the Step 4 analysis.

##### TASK

Your task is to compare within-subject EEG groups, and to decide, for each subject separately, which group is “dreaming”, and which is “dreamless sleep”. At this stage, 3 vs. 3 EEGs can be grouped together and within-subject averaging can be made. Yet, between-subject comparisons are still impossible.

##### RESULTS

At least 14 EEG pairs of samples out of 9 should be correctly identified in order to succeed above chance level. Also, each research group is expected to provide methodological rationale, on what EEG basis the decision was made, that is, which features in the EEG predict dreaming or non-dreaming.

Feedback will be given about overall success rate only, not about correct classification of individual groups.

#### Step 5: *Conditions* of data set are provided

##### DATA

Finally, all data will be classified between-subjects into 2 conditions with 27 EEG cases per each (see Fig. 5).

**Figure 5.**
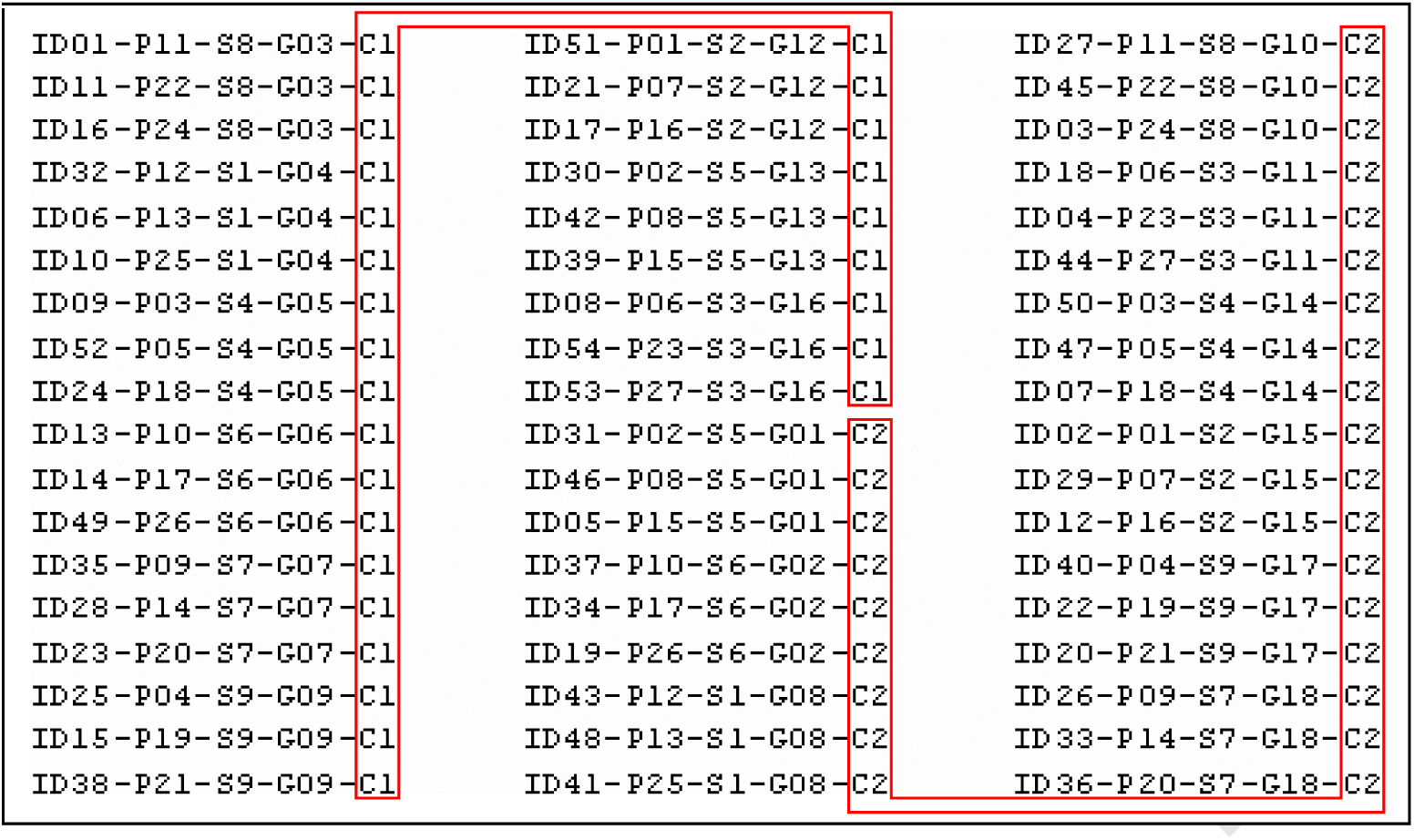
Example of 54 coded files names for the Step 5 analysis.

##### TASK

Your task will be to decide, which one of the two conditions is “dreaming” and which is “dreamless sleep”. All possible analysis can be made at this stage, as both within- and between-subjects averaging is possible.

##### RESULTS

Each research group will report findings of comparison between these two groups. Unfortunately, correct identification of the groups won’t be accepted as success above chance level anymore at this stage (because there is 50% chance to get it right by guessing).

## Supplementary Document 5

### Themed Feature Set Candidates for Step 1 Blind Classification

The Analysis Team constructed eight sets of features, which were themed around various analyses of time series data. The sets were called: *Power*, *PowerFine*, *ACC*, *PermEn*, *ApEn*, *Siclari*, *EogRms* and *EmgRms*. Their constructions are explained below. As described in the main text, the team eventually chose the *PowerFine* features for Step 1 of blind classification due to its high temporal consistency. See Table S5.1 for a summary.

**Table S5.1.**
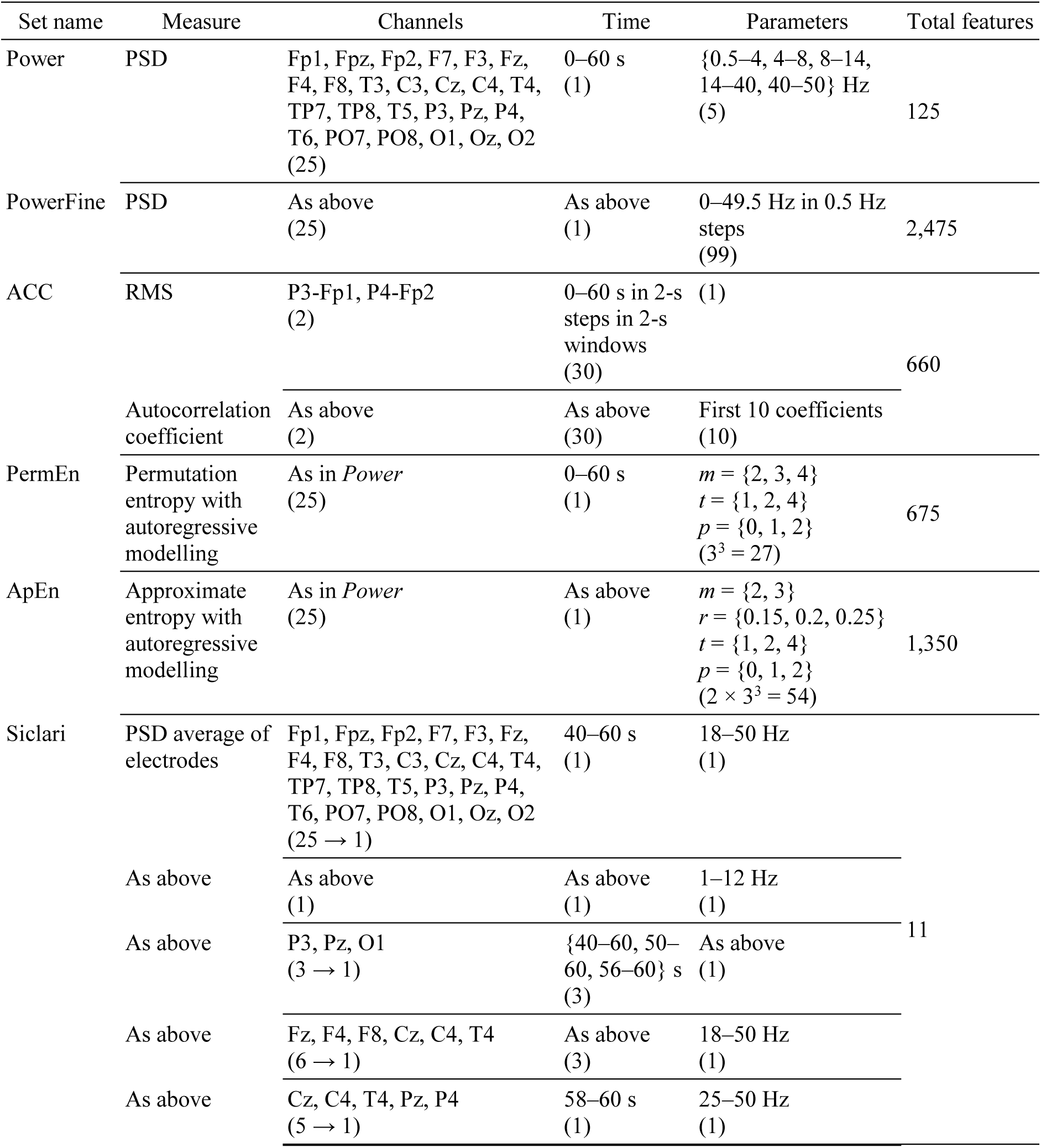
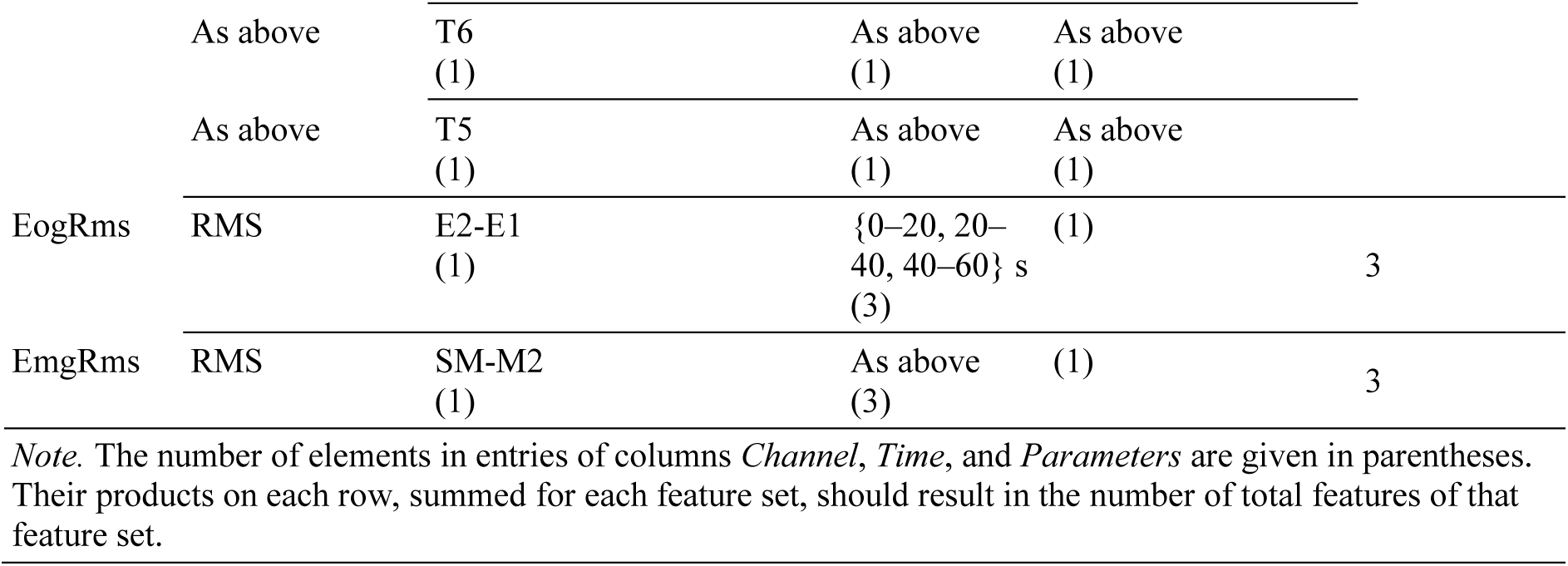
Candidate feature sets of Step 1

### Power

The Analysis Team constructed the *Power* feature set to provide information about both brain activity levels and locations. The set consisted of PSD estimates for each EEG electrode. They evaluated PSDs over the whole 60-s duration in five frequency bands corresponding to delta (0.5–4 Hz), theta (4–8 Hz), alpha (8–14 Hz), beta (14–40 Hz) and gamma (40–50 Hz) wave activity (Noachtar et al., 1999). The electrodes used were a subset of the 10-20 and 10-10 system electrodes (Chatrian, Lettich, & Nelson, 1985; Klem, Luders, Jasper, & Elger, 1999) totalling 25 in number: Fp1, Fpz, Fp2, F7, F3, Fz, F4, F8, T3, C3, Cz, C4, T4, TP7, TP8, T5, P3, Pz, P4, T6, PO7, PO8, O1, Oz and O2. The feature set in total had 125 features (5 frequency bands × 25 channels) for each case.

The Analysis Team preprocessed the EEG time series data offline using the EEGLAB Matlab toolbox (version 13.5.4b; Delorme & Makeig, 2004). From the raw EEG recordings, they detrended the signals and downsampled them from 2000 Hz sampling rate to 100 Hz. They then applied a finite impulse response high-pass filter, windowed using a Hann window of 31 s, at the cutoff frequency of 0.1 Hz. Finally, they re-referenced the channels to the average common electrode. This preprocessed data is also reused in the methods for other feature sets.

PSDs for each electrode were estimated using a modified Welch’s method (Welch, 1967) with time segments of 6.2 s overlapping by 80%, windowed by Hann windows. The Analysis Team performed fast Fourier transform on each window. The power of a frequency band was taken as the mean power (not log-transformed) across frequency bins, from and including the lower limit of the band, up to and exclusive of the upper limit. The Analysis Team modified Welch’s method by trimming off time segments with values in both the upper and lower 5% range prior to taking the mean power of the remaining time segments for each band.

### PowerFine

The *PowerFine* feature set was identical the *Power* feature set, except that the PSDs were calculated for much finer-resolution, more uniformly-spaced frequency bins. This was chosen to offer more complete frequency information. The frequency edges of the bins were from 0 to 49.5 Hz in 0.5 Hz steps. PSDs for each electrode were estimated using the modified Welch’s method with time segments of 9.3 s overlapping by 80%, windowed by Hann windows. The feature set in total had 2,475 features (25 channels × 99 frequencies) extracted for each observation.

### ACC

This feature set’s name was abbreviated from “autocorrelation coefficients”, and consisted of features replicated from a study to assess anaesthetic depth following machine learning through cluster analysis (Thomsen et al., 1991). The performance of their method appeared superior to other single parametric measures of spectral distribution (i.e., median and spectral edge frequencies) suggested for assessing anaesthetic depth at the time. Their features were based on autoregressive modelling of EEG signals in 2-s time windows, taking the form of 10 normalised autocorrelation coefficients and 1 root-mean-square measure (RMS).

To replicate this feature set, the Analysis Team utilised the preprocessed (high passed and re-referenced) EEG data performed for the *Power* feature set. From the preprocessed EEG, two bipolar–re-referenced channels were extracted to summarise the activity of the two hemispheres of the brain: electrodes P3-Fp1 and P4-Fp2. They then further filtered those channels, in following with Thomsen’s method, with a pre-emphasising first order high-pass filter at 4.2 Hz, and a fourth order low-pass filter at 25 Hz using Matlab’s Butterworth filter design tools. For each channel, the Analysis Team segmented the signal into 30 × 2-s time segments, and calculated the RMS and first 10 normalised sample autocorrelation coefficient lags of each segment. The feature set in total had 660 features (2 bipolar channels × 30 time segments × 11 coefficients) extracted for each observation.

### PermEn and ApEn

The Analysis Team explored information theory measures of time series data in the form of entropy: specifically through permutation entropy (Bandt & Pompe, 2002) in the *PermEn* feature set, and approximate entropy (Pincus, Gladstone, & Ehrenkranz, 1991) in the *ApEn* feature set. Entropy in signals is a measure of the complexity of a system from which the signals were taken. In the context of consciousness research, entropy has been proposed as a way to monitor loss of consciousness via anaesthetic depth (Bein, 2006; Liang et al., 2015).

For the *PermEn* feature set, the Analysis Team used a Matlab implementation by Ouyang (2012) to compute measures of permutation entropy. Specifically, for each channel of EEG recording (after re-referencing followed by high-pass filtering, as in *Power* feature set), they extracted permutation entropy for all 27 combinations of three parameters of the analysis: the embedding dimension *m* (2, 3 or 4), the downsampling time delay *t* (1, 2 or 4), and autoregressive order *p* (0, 1 or 2). Parameters *m* and *t* were direct inputs for the permutation entropy function. The Analysis Team varied *m* no higher than 4 so as to conserve computation time. Before computing permutation entropies, they performed autoregressive modelling of order *p* on single EEG signals, and then subtracted the modelled time course from the EEG signals. This was done to reveal the evolution of a time series that may not be linearly dependent on its immediate past states, which may closer reflect signals related to consciousness. The Analysis Team carried out autoregression in Matlab with the function provided by the System Identification Toolbox, using the default forward-backward approach to the least-squares autoregressive fitting algorithm. The feature set in total had 675 features (25 electrodes × 3 embedding dimensions × 3 downsampling delays × 3 autoregressive orders) extracted for each case.

For the *ApEn* feature set, the Analysis Team used a Matlab function implemented by Lee (2012) to calculate approximate entropy. Similarly to *ApEn*, for each preprocessed channel of EEG recording, they extracted approximate entropy for all 54 combinations of four parameters: the embedding dimension *m* (2 or 3), the filter tolerance *r* (0.15, 0.2 or 0.25; relative to each signal’s sample standard deviation), the downsampling time delay *t* (1, 2 or 4), and autoregressive order *p* (0, 1 or 2). The values for *m* were typical choices by Pincus et al (1991). Like in *PermEn* the Analysis Team subtracted the *p*^th^-order autoregressed time course before computing approximate entropy. The feature set in total had 1,350 features (25 channels × 2 embedding dimensions × 3 filter tolerances × 3 downsampling delays × 3 autoregressive orders) extracted for each observation.

### Siclari

The extraction of the *Siclari* feature set was designed to approximate the features described by Siclari, LaRocque, Bernardi, Postle, & Tononi (2014) in their preprint manuscript of a study later published in Nature Neuroscience (Siclari et al., 2017). They found a number of significant differences in low- and high-frequency activity of specific brain regions, as recorded with high-density EEG (256 channels), correlating with conscious experience during sleep. Due to the technical differences between our protocols, the *Siclari* feature set here does not constitute a direct replication. We note the following differences in our protocols. (a) We had considerably fewer channels recorded (25 vs. 256). Owing to this, (b) the Analysis Team deferred from performing source localization, as its reliability is lowered by decreasing the number of channels. For the same reason, (c) they did not perform independent component analysis for the removal of ocular, muscular and cardiac artefacts; however, cases were already chosen to have a minimum of such artefacts.

The *Siclari* feature set consisted of 11 of these differences: the average power across all electrodes at low- (1–12 Hz) and high-frequencies (18–50 Hz); low-frequency power in parieto-occipital channels over the 20, 10 and 4 s till awakening; high-frequency power in frontal channels over the final 20, 10 and 4 s till awakening; and high-frequency (25–50 Hz) power over the final 2 s preceding awakening at regions correlated with conscious experience of either spatial setting, movement, or speech.

The Analysis Team utilised the preprocessed EEG data performed for the *Power* feature set and performed further operations in order to derive a scalp current source density estimate (CSD) of the EEG signal. They calculated the CSD using Perrin’s method of taking the Laplacian of spherical splines (Perrin et al., 1989; Perrin, Pernier, Bertrand, & Echallier, 1990), as implemented for Matlab in CSD Toolbox (Kayser, 2010; Kayser & Tenke, 2006a, 2006b). The parameters for this method were based on those recommended by Kayser & Tenke (2015): a spline flexibility of 4, and smoothing constant of 10^-5^. The Analysis Team assumed a head radius of 8.9 cm, and thus produced CSDs for each of the 25 original electrode locations for each case.

PSDs for each electrode were estimated following Siclari et al.’s procedure: using Welch’s method with time segments of 2 s overlapping by 50%, windowed by Hamming windows. They performed fast Fourier transform on each window. The power of a frequency band was taken as the mean (not log-transformed) power across frequency bins, inclusive of the band’s specified edges.

The 11 features were extracted as follows. Two features of power for low and high frequencies respectively were taken as means across all electrode locations in the bands 1–12 Hz and 18–50 Hz over the final 20 s preceding awakening. Three features of low-frequency, average parieto-occipital power were taken across electrodes P3, Pz and O1 in the band 1–12 Hz over the final 20, 10 and 4 s till awakening. Three features of high-frequency, average frontal power were taken across electrodes Fz, F4, F8, Cz, C4 and T4 in the band 18–50 Hz over the final 20, 10 and 4 s till awakening. Three features for dream contents were taken: spatial setting content power as the average power over electrodes Cz, C4, T4, Pz and P4 in the band 25–50 Hz for the final 2 s till awakening; movement content power as that of electrode T6; and speech content power as that of electrode T5.

### EogRms and EmgRms

The *EogRms* and *EmgRms* feature sets contained information about eye movement and muscle tone, measured as RMS, over three time segments.

For *EogRms*, the Analysis Team used electrooculograms, taken as the bipolar re-referenced channel E2-E1, resampled to a rate of 60 Hz from 2000 Hz. They applied a first-order Butterworth high-pass filter with a cutoff frequency of 0.5 Hz, to remove particularly slow voltage changes and not saturate the RMS calculation. Three RMS features were calculated for equally divided time segments: 0–20, 20–40 and 40–60 s from awakening.

For *EmgRms*, the Analysis Team used electromyograms, taken as the bipolar re-referenced channel SM-M2, retaining its 2000 Hz sampling rate and online 5–500 Hz bandpass. Three RMS features were calculated for equally divided time segments as in *EogRms*.

## Supplementary Document 6

## Feature Sets and Sub-Clustering Used in Steps 2–5 of Blind Classification

The Analysis Team tailored and carried out unique procedures for each step of blind classification. A summary of their extracted features for each step is given in Table S6.1, and details are described thereafter.

**Table S6.1.**
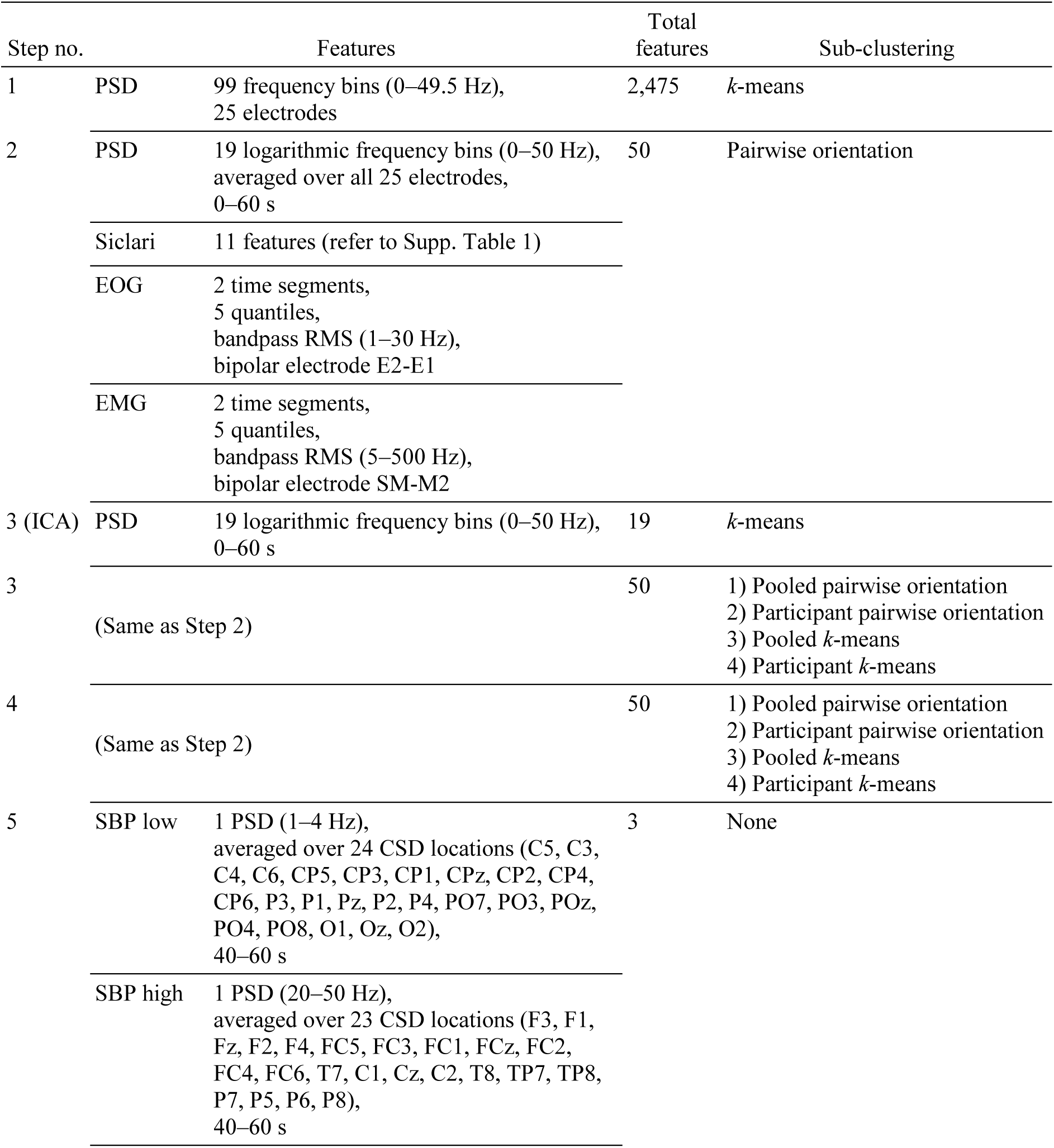
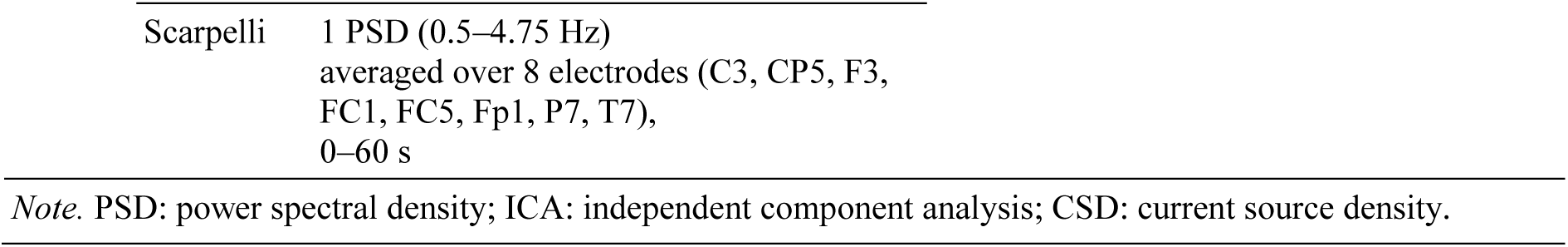
Summary of feature sets and sub-clustering used in the blind classification experiment

| Step no. | Features |  | Total features | Sub-clustering |
| --- | --- | --- | --- | --- |
| 1 | PSD | 99 frequency bins (0–49.5 Hz),<br>25 electrodes | 2,475 | *k*-means |
| 2 | PSD | 19 logarithmic frequency bins (0–50 Hz),<br>averaged over all 25 electrodes,<br>0–60 s | 50 | Pairwise orientation |
|  | Siclari | 11 features (refer to Supp. Table 1) |  |  |
|  | EOG | 2 time segments,<br>5 quantiles,<br>bandpass RMS (1–30 Hz),<br>bipolar electrode E2-E1 |  |  |
|  | EMG | 2 time segments,<br>5 quantiles,<br>bandpass RMS (5–500 Hz),<br>bipolar electrode SM-M2 |  |  |
| 3 (ICA) | PSD | 19 logarithmic frequency bins (0–50 Hz),<br>0–60 s | 19 | *k*-means |
| 3 | (Same as Step 2) |  | 50 | 1) Pooled pairwise orientation<br>2) Participant pairwise orientation<br>3) Pooled *k*-means<br>4) Participant *k*-means |
| 4 | (Same as Step 2) |  | 50 | 1) Pooled pairwise orientation<br>2) Participant pairwise orientation<br>3) Pooled *k*-means<br>4) Participant *k*-means |
| 5 | SBP low | 1 PSD (1–4 Hz),<br>averaged over 24 CSD locations (C5, C3,<br>C4, C6, CP5, CP3, CP1, CPz, CP2, CP4,<br>CP6, P3, P1, Pz, P2, P4, PO7, PO3, POz,<br>PO4, PO8, O1, Oz, O2),<br>40–60 s | 3 | None |
|  | SBP high | 1 PSD (20–50 Hz),<br>averaged over 23 CSD locations (F3, F1,<br>Fz, F2, F4, FC5, FC3, FC1, FCz, FC2,<br>FC4, FC6, T7, C1, Cz, C2, T8, TP7, TP8,<br>P7, P5, P6, P8),<br>40–60 s |  |  |

|  |  |
| --- | --- |
| Scarpelli | 1 PSD (0.5–4.75 Hz)<br>averaged over 8 electrodes (C3, CP5, F3,<br>FC1, FC5, Fp1, P7, T7),<br>0–60 s |

*Note.* PSD: power spectral density; ICA: independent component analysis; CSD: current source density.

### Step 2

The *PowerFine* feature set used in Step 1 of blind classification consisted of 2,475 features (25 EEG channels × 99 frequencies) that measured only EEG PSDs. The resulting classification accuracy was near chance level. By Step 2, the Analysis team suspected that the poor performance might have resulted partly from the large number of features that they used, causing overfitting of data (Domingos, 2012). They also suspected that their clustering algorithm might have performed better with a more encompassing feature set than one that looked only at EEG. The Analysis Team therefore decided to change the feature set for Step 2 to be more concise and to include information from EMG and EOG (different to that from Step 1). Summarily, they reduced the number of EEG features from Step 1 by taking the average PSD over all 25 electrodes and reducing the number of frequency bins to 19; to avoid losing all spatially specific EEG information, they added PSD estimates for localised brain areas based on the Siclari et al. (2014) report (identical to the Siclari feature set in Supplementary Document 5). In total, the Analysis Team used 50 features for Step 2 (19 from average EEG, 10 from EMG, 10 from EOG, and 11 from Siclari EEG)—more than a 100-fold reduction from Step 1.

For 19 features of average EEG power, the Analysis Team evaluated PSDs for each electrode in 18 frequency bins with edges logarithmically spaced between 1.3^-3^ and 1.3^15^ Hz (approximately 0.46–51 Hz), and a lowermost bin with edges at 0 and 1.3^-3^ Hz. These were evaluated in a way similar to the *Power* feature set in Step 1 (see Supplementary Document 5), with one difference being that the time segments used for Welch’s method have lengths of 9.3 s instead of 6.2 s.

Eleven more features were taken directly from the *Siclari* feature set from Supplementary Document 5. Lastly, they extracted 10 features each for EOG and EMG modalities. The data was preprocessed similarly to *EogRms* and *EmgRms* as described in Supplementary Document 5, but for 2 time segments each instead of 10. Thus, for each of these 30-s segments of EMG or EOG, they computed the RMS values of all consecutive 1-s time windows, and took the 0^th^, 25^th^, 50^th^, 75^th^, and 100^th^ percentiles of those as features.

### Step 3

The analysis in Step 3 followed two stages. In the first stage, to take advantage of the revealed participant grouping, the Analysis Team used independent component analysis (ICA) to remove specific components that were likely to affect incorrect clustering results. They considered a component to cluster incorrectly if any pair of cases were to be clustered together rather than apart. The first stage involved extracting the independent components, testing them for incorrect clustering, removing components that were significantly incorrect, and then recomposing the remaining components back into cases for feature extraction. The second stage consisted of combination clustering using a more elaborate sub-clustering procedure than previous Steps. Both stages will now be described.

#### ICA

The ICA cleaning stage was performed on each participant-group of cases. See Fig. S6.1 for a schematic overview.

**Figure S6.1.**
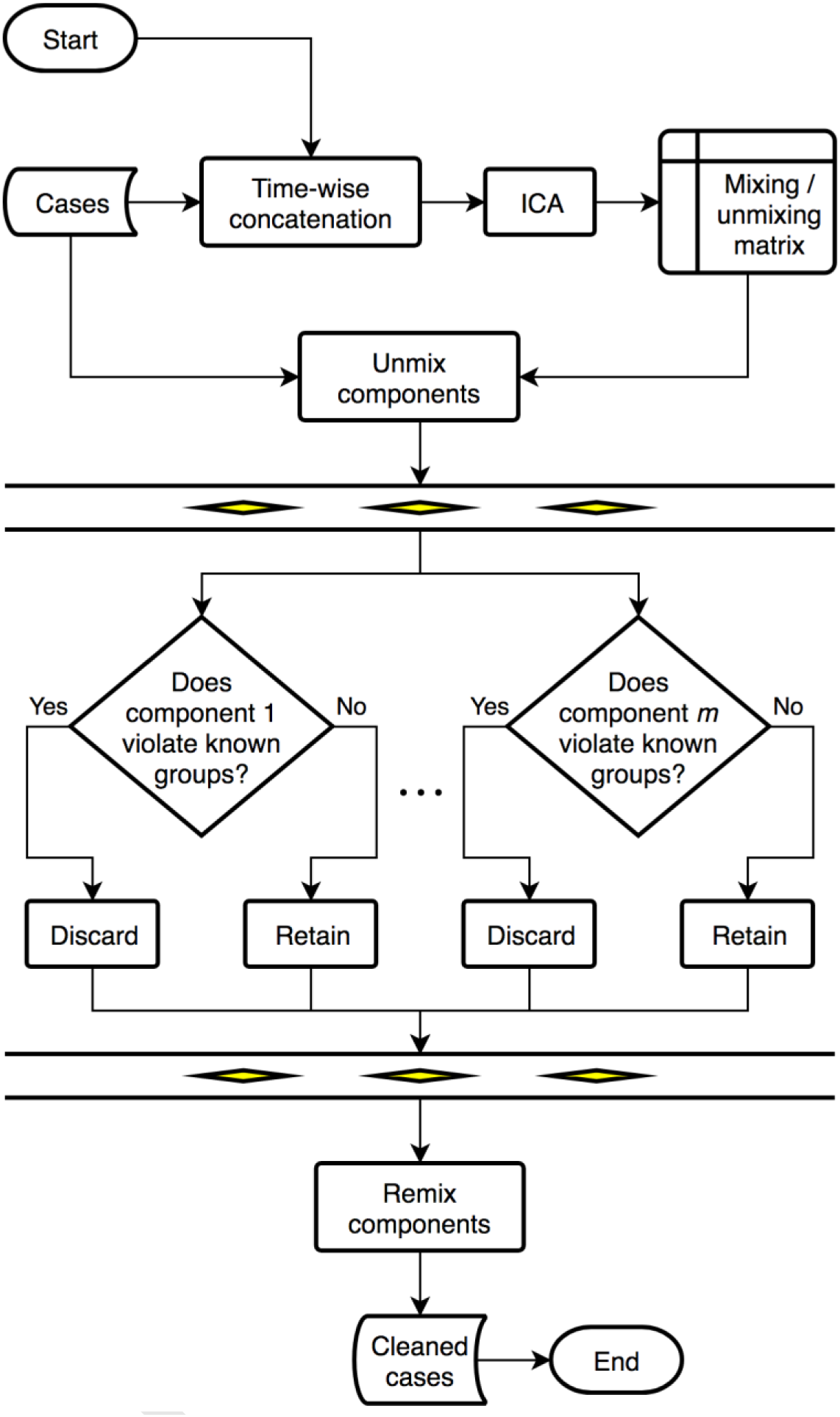
Flowchart overview of ICA cleaning. The procedure is performed for each participant-group of cases.

Using the software package *FastICA for Matlab 7.x and 6.x* (version 2.5; Gävert, Hurri, Särelä, & Hyvärinen, 2005), the Analysis Team applied ICA to the combined EEG and EOG time series, concatenating in time across all six cases for a given participant. Independent components were computed using the FastICA “symmetric” approach with a “tanh” nonlinearity function (Rogasch et al., 2014).

At this step of blind classification, case pairing information (that each pair came from the same participant and had different dream report conditions) was already known. Therefore, each of the 26 independent components could be used to cluster the cases of a single participant, and components that reliably clustered contrary to pairing information (i.e., components that represented dissimilar cases as similar) could be identified and removed from the original case recording. The Analysis Team identified these problem components by observing whether they reliably clustered cases of the same pair together rather than apart. To do this, they used a feature set consisting of spectral power in 19 frequency bins, calculated in identical fashion to that in Step 2. Ensemble clustering with modifications was carried out on this feature set extracted for each independent component. Due to the small sample size (three cases) of each participant-group, the Analysis Team tried to avoid overfitting by sub-clustering in up to three dimensions with *k* = 3, and not weighting sub-cluster results by their quality of clustering. They also replaced the agglomerative clustering step of combination clustering with one that used an exhaustive cluster configuration search on the co-association similarity matrix, made feasible due to the small sample size per participant-group.

The search objective operationalised by finding the clustering outcome that maximised overall cluster quality as measured by the mean silhouette value (Rousseeuw, 1987). Silhouette values quantify the cohesion of individual members of a cluster and their separation from other clusters, as expressed in the following equation:

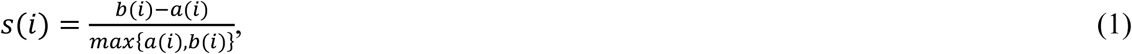

where *a*(*i*) is the average dissimilarity (i.e., distance) of member *i* to all other members of its cluster, and *b*(*i*) is the average dissimilarity of member *i* to all members of the next nearest cluster.

For the exhaustive search, the Analysis Team restricted the set of possible clusterings to search within by considering only configurations consisting of four clusters: one cluster to represent three cases from one dream report condition group, and the remaining other three clusters to each represent one case from the other dream report condition group. This restriction was imposed to accommodate the possibility of one dream report condition having higher variance cases than the other. There were 20 such configurations (6 choose 3).

An independent component was considered to cluster incorrectly when, after clustering on its extracted feature set, the larger cluster did not consist of exactly one case from each pair. The statistical significance of this effect was measured by bootstrapping the sub-clustering results on which the evidence accumulation clustering was performed. The Analysis Team bootstrapped 1,000 similarity matrices by randomly sampling from the pool of all sub-clustering results with replacement. Formally, they tested the hypothesis that the independent component being considered clustered incorrectly with respect to dream report condition by assuming the null hypothesis that they clustered relevantly to condition. The Analysis Team would reject this null hypothesis if the proportion of bootstrapped clusterings, in which the larger cluster consisted of exactly one case from each pair, was lower than 5%. Significantly incorrect independent components by this criterion were subsequently removed. They then recomposed the EEG and EOG traces from the remaining independent components. On average, 14 components were retained, with a range of 5 to 23.

##### Sub-clustering

A new set of sub-clustering procedures was devised to take into account the newly revealed participant labels. The Analysis Team could remove unwanted participant effects by sub-clustering cases within each participant; however, this would prevent the analysis from finding similarities amongst cases between participants. The team also suspected that the method of sub-clustering in Step 2—dividing case pairs along the mean orientation of the difference of pairs—may have been discarding other useful information about in the cases’ actual positions in feature space, as was possible with *k*-means clustering. Therefore, they intended to devise a sub-clustering procedure that additionally retained clustering information from unconstrained case features and from the whole pool of participants.

In fact, the Analysis Team sub-clustered the cases in four different ways and combined the results. In two of the ways, similar to Step 2, they sub-clustered cases in a pairwise manner: firstly with respect to the mean orientation amongst all pairs, and secondly with respect to the mean orientation of each pair’s own participant. In the other two ways, they took the method of Step 1’s *k*-means sub-clustering and sub-clustered the unpaired cases: firstly amongst all cases, and secondly amongst each participant-group of cases. Therefore, there were four ways which they performed sub-clustering by. The scheme is illustrated in Figure S6.2. After correcting for their average pairwise similarity distance, the Analysis Team averaged those results, thereby performing evidence accumulation, to obtain the final co-association similarity matrix. For per-participant sub-clusterings, they limited the maximum number of features in a combination to 2 due to the effective reduction in data size. What follows are the details for each of the four ways of sub-clustering.

**Figure S6.2.**
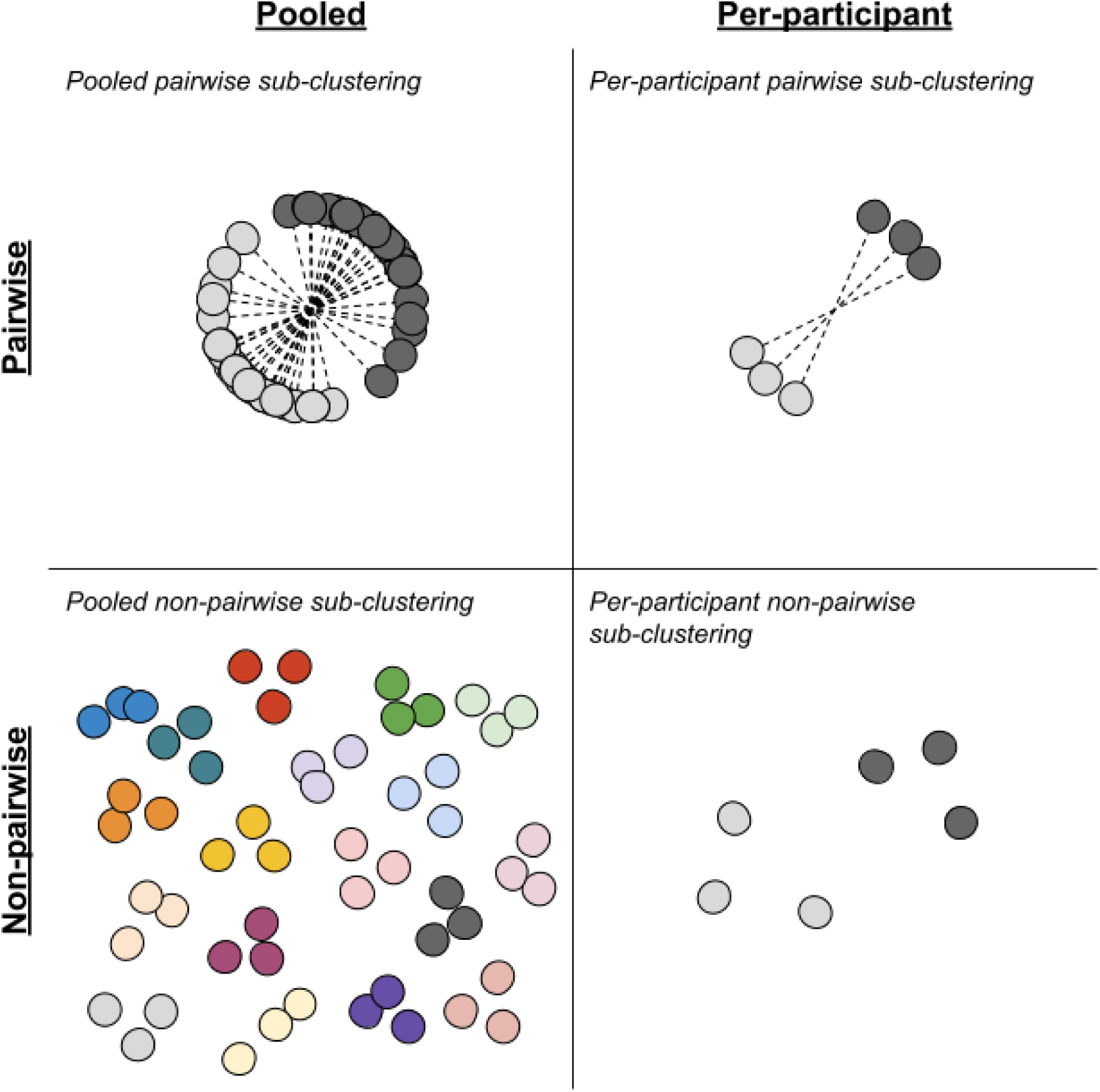
Four Step 3 sub-clustering schemes. Each circle corresponds to a case in feature space; each dashed line corresponds to a case pair or difference vector. Colours indicate different sub-clusters that cases belong to. Non-pairwise sub-clustering was implemented as *k*-means clustering.

##### Pooled pairwise sub-clustering

For sub-clustering cases in the pooled pairwise manner, the Analysis Team dichotomously grouped each case of every pair with respect to the average pair difference vector orientation across all participants, as done in Step 2. For each sub-cluster’s resulting co-association similarity matrix, prior to evidence accumulation, they weighted the matrix such that they took on a mean co-association distance equal to their goodness of clustering value—being 1 minus the mean cosine similarity of the difference vectors.

##### Pooled non-pairwise sub-clustering

For sub-clustering cases in the pooled non-pairwise manner, the Analysis Team performed *k*-means clustering over all cases with *k* = 18 (the number of participant-condition groups in our data), after subtracting the means of each participant. For each sub-cluster’s resulting co-association similarity matrix, prior to evidence accumulation, they forced any incorrectly co-associated pairs of cases to be recorded as not associated, and then weighted the matrix to take on a mean co-association distance equal to their mean silhouette value.

##### Per-participant pairwise sub-clustering

For sub-clustering cases in the per-participant pairwise manner, the Analysis Team dichotomously grouped each case of every pair per participant with respect to the participant’s average pair difference vector orientation. They weighted each sub-cluster’s co-association similarity matrix prior to evidence accumulation just like for pooled pairwise sub-clustering. They then combined the results from all individual participants into the co-association similarity matrix for all cases, where the uncalculated pairwise similarity distances between different participants were simply set to the average of the calculated similarity distances.

##### Per-participant non-pairwise sub-clustering

For sub-clustering cases in the per-participant non-pairwise manner, the Analysis Team performed *k*-means clustering on all cases per participant with *k* = 4 (which accommodates for one low-variance and one high-variance dream report condition group for each participant, where the high-variance group consisted of three single-membered clusters). For each sub-cluster’s resulting co-association similarity matrix, prior to evidence accumulation, they forced any incorrectly co-associated pairs of cases to be recorded as not associated, then weighted the matrix prior to evidence accumulation just like for per-participant non-pairwise sub-clustering. They then combined the results of all individual participants into the co-association similarity matrix for all cases, just like for per-participant pairwise sub-clustering.

### Step 4

Like in Step 3, ICA was performed with the intention to remove specific independent components that were likely to affect incorrect clustering results. Specifically, the Analysis Team tested components for clustering contrary to the revealed participant-condition grouping information in addition to pairing information.

They slightly altered the statistical procedure for testing the significance of bad independent components. Instead of rejecting the null hypothesis if the proportion of valid bootstrapped clusterings was lower than 5%, they calculated a different threshold for each component tested using the Benjamini–Hochberg–Yekutieli procedure (Benjamini & Yekutieli, 2001) to control the false discovery rate to α = .05. On average, four components were retained, with a range of one to six.

For the proper combination clustering stage on the recomposed case–averaged participant-condition groups, the Analysis Team sub-clustered these groups as nine pair difference vectors in combinations of up to eight features. They used the same sub-clustering procedure as that of Step 2 on these difference vectors.

### Step 5

In Step 5, the Analysis team extracted features based on significant differences reported by Siclari et al. (2017) and Scarpelli et al. (2017). Two features were the low frequency and high frequency power, which was named respectively *SBP low* and *SBP high* (SBP for the initials of the first three authors of the Siclari paper), and one feature was of the low frequency activity reported by Scarpelli et al., which was named *Scarpelli*.

To replicate the SBP features, they started by downsampling the raw 2000 Hz EEG case recordings to 500 Hz sampling rate and applying a band-pass filter for frequencies 1–50 Hz. The EEG montage was then re-referenced to the common average electrode. They next calculated the CSD using Perrin’s method, as described in Supplementary Document 5 for the *Siclari* feature set. Electrodes for *SBP low* were C5, C3, C4, C6, CP5, CP3, CP1, CPz, CP2, CP4, CP6, P3, P1, Pz, P2, P4, PO7, PO3, POz, PO4, PO8, O1, Oz and O2; for *SBP high* were electrodes F3, F1, Fz, F2, F4, FC5, FC3, FC1, FCz, FC2, FC4, FC6, T7, C1, Cz, C2, T8, TP7, TP8, P7, P5, P6 and P8. Electrodes were interpolated where missing using spherical splines. The Analysis Team next estimated the PSDs of each electrode for the 20-s duration until awakening using Welch’s method with 2-s Hamming windows and 50% overlap. Finally, for *SBP low*, they took the average power across all its electrodes between 1 and 4 Hz; and for *SBP high*, they took the average power across all its electrodes between 20 and 50 Hz.

To replicate Scarpelli’s feature, the Analysis Team started by resampling the raw EEG case recordings down to 250 Hz and applying a band filter for frequencies 0.5–30 Hz. They then retained or interpolated electrodes C3, CP5, F3, FC1, FC5, Fp1, P7 and T7 using the spherical spline method. They next estimated PSDs of each electrode, using Welch’s method with 4-s rectangular windows and no overlap, for frequencies 0.5 to 4.75 Hz in 0.25 Hz steps. Finally, they calculated the average power across the above-mentioned 8 electrodes and frequencies, and took its natural logarithm as the feature.

## Supplementary Document 7

### Test Decoding of Electrocorticograms

The Analysis Team tested the decoder, used in the Dream Catcher post hoc analysis, on a different set of electrophysiological recordings with known representation of visual stimulus– evoked information. They demonstrated its utility in decoding this embedded information with a high success rate.

#### Method

##### Data

The experimental setup and collection of data used here have been described in previously published studies; please refer to Baroni et al. (2017) and Haun et al. (2017). The Analysis Team used data consisting of electrocorticogram evoked potentials from epilepsy patient 153, taken during the period of epilepsy monitoring after electrode implantation. Out of a grid electrode array installed over the right temporal lobe, they used recordings from two bipolar re-referenced channels, found over the ventral fusiform area, that were sensitive to faces. No recordings were performed within 48 hours of a major seizure. Visual stimuli—consisting of upright faces, upside-down faces, houses, line drawings of tools, and Mondrian patterns— were presented in a continuous cycle while the participant fixated on a cross at the centre of the display.

The Analysis Team arranged the data recordings into a set of two groups for four separate decoding experiments—each group was composed of a set of 27 evoked potentials to either face or non-face stimuli, and from either one or two electrodes, as shown in Table S7.1.

**Table S7.1.** Data sets

| Data set | Stimuli | Electrodes | Timing (ms) |
| --- | --- | --- | --- |
| A | Face vs. non-face | 2 intermixed | 100–300 |
| B | Face vs. non-face | 2 intermixed | 400–600 |
| C | Face vs. Mondrian | Single | 100–300 |
| D | Face vs. Mondrian | 2 intermixed | 400–600 |

*Note.* Face stimuli consisted of both upright and inverted faces. Non-face stimuli consisted of houses, tools, and Mondrian images. Timing is relative to stimulus onset.

For each evoked potential, they extracted a set of 19 power spectral density (PSD) features in 19 logarithmically spaced frequency bins between 0 Hz and the Nyquist frequency, based on the 19 EEG PSD features used in Step 2 of the Dream Catcher blind classification experiments (see Supplementary Document 6). They calculated PSDs using the multitaper method (Thomson, 1982) with five Slepian tapers before binning. The Analysis Team evaluated their decoder by decoding for stimulus category in each data set.

##### Decoder

The Analysis Team used a support vector machine (SVM; Cortes & Vapnik, 1995) for decoding. Specifically, they employed the LIBSVM implementation for Matlab (version 3.18; Chang & Lin, 2011, 2014). They used the nu-SVC model, with a Gaussian kernel type, the regularisation parameter set to the default *nu* = 0.5, and the model set to output probability estimates. They estimated decoding accuracy using leave-one-out cross-validation on the data as partitioned into pairs of face-non-face evoked potentials. For each fold of cross-validation, the Analysis Team classified the pairs dichotomously based on their relative probability estimate outputs. They preprocessed the power features by taking their natural logarithm, and standardising them across all features by Studentisation.

#### Results

The supervised SVM decoder generally performed well for all data sets (Table S7.2). The average accuracy across all sets was 89%, with the lowest being 74% for Set B. Their 95% confidence intervals, estimated using the Clopper-Pearson method (Clopper & Pearson, 1934), were all greater than 50%, indicating statistically significant performance in all sets.

**Table S7.2.** Decoding results

Decoding results
| Data set | Cross-validation accuracy |  |  |
| --- | --- | --- | --- |
|  | *n* correct<br>(out of 27) | % |  |
|  |  | Average | 95% CI |
| A | 25 | 93 | [76, 99] |
| B | 20 | 74 | [54, 89] |
| C | 27 | 100 | [87, 100] |
| D | 24 | 89 | [71, 98] |
| *Note.* CI: confidence interval. All values were statistically significant. |  |  |  |

The Analysis Team achieved the highest accuracy in data set C, in which the evoked potentials were not of mixed electrodes (one-tailed Fisher’s exact test against all other sets collapsed, *p* = .025). This suggested that the inclusion of a second, unrelated factor negatively impacted the decoding performance, but the decoder was resilient enough to preserve significant discriminability.

